# Donor Age Impairs Vasculogenic Potential of hiPSC-Derived Endothelial Progenitors

**DOI:** 10.1101/2025.06.24.661422

**Authors:** Bryce Larsen, Cody Callahan, Arshitha Rayanki, Sara Faulkner, Janet Zoldan

## Abstract

**Background:** Human induced pluripotent stem cells (hiPSCs) hold promise for vascular regeneration, but preliminary research often relies on neonatal donors, whereas clinical applications will use cells derived from older adults. Although the impact of donor age on reprogramming efficiency has been studied, its effect on the functionality of hiPSC-derived endothelial progenitors (hiPSC-EPs) remains unclear. This question is the focus of the current study.

**Methods and Results:** We derived EPs from iPSCs sourced from three neonatal donors (ND) and three mature donors (MD) matched 1:1 for sex and somatic cell origin. We assessed their functional, epigenetic, and transcriptomic characteristics. Despite higher CD34⁺ yields from MD-iPSCs, MD-hiPSC-EPs formed poorly interconnected and non-lumenized vascular structures in 3D hydrogels, compared to neonatal donor (ND) lines. In 2D culture, MD-hiPSC-EPs exhibited reduced cell density and aberrant VE-Cadherin localization. DNA methylation analysis revealed that somatic cell origin was the dominant driver of variance, but consistent differences in methylation of mesoderm commitment, angiogenesis, ECM remodeling, and cytoskeleton-related genes were observed between age groups. Epigenetic age prediction showed MD-hiPSC-EPs had more developmentally advanced signatures, potentially explaining their shift away from vasculogenic competence. Our RNA-sequencing findings confirm trends seen in the DNA methylation data and show differential expression of pathways linked to mitochondrial regulation and nitric oxide signaling.

**Conclusions:** Donor age significantly alters the vasculogenic function of hiPSC-EPs. These findings underscore the necessity of donor-specific considerations in hiPSC-based vascular engineering and highlight potential barriers to translating hiPSC-derived therapeutics into older patient populations.

## Introduction

Cardiovascular diseases remain the leading cause of mortality worldwide and are especially prevalent in the United States (1,2). While several risk factors are lifestyle-dependent, aging stands out as a significant and unavoidable contributor (3). Some of the prominent age-related changes driving cardiovascular disease include a decline in vascular density and function. In older adults, studies show rarefaction of the microvasculature in multiple organs, including the heart (4). This loss, combined with a diminished regenerative capacity in aged tissues, is a critical challenge for current therapies that do not restore or replace damaged and lost vasculature.

Given these challenges, engineering and replacing aged or diseased tissue presents a promising strategy for addressing cardiovascular disease in older adults (5). One such approach involves generating new vascular tissue using human induced pluripotent stem cells (hiPSCs) (6). Derived from somatic cells, hiPSCs are patient-specific and possess the ability to self-renew and differentiate into a wide array of cell types, including vascular cells (7). This differentiation capacity makes them a valuable tool for studying and potentially regenerating damaged vasculature. Their patient specificity also opens avenues for personalized preclinical models and therapies.

While this patient specificity offers significant advantages, it also introduces variability based on the characteristics of the donor cell. For instance, somatic cell origin has been shown to influence hiPSCs’ differentiation propensity to endothelial progenitor cells (hiPSC-EPs) (8). However, the impact of hiPSC donor age on hiPSC-EP functionality remains largely unexamined. Most existing work on the impact of donor age on differentiated hiPSCs has primarily focused on differentiation propensity, without quantitatively investigating how age-related factors affect downstream functionality. (9). This gap in understanding how donor background impacts functionality presents a critical barrier to translating hiPSC-based vascular therapies into clinical settings.

Furthermore, current hiPSC-derived vascular engineering heavily relies on somatic cells obtained from newborn or fetal donors (10–15). Although these neonate donor-hiPSCs (ND-hiPSCs) are ideal in a laboratory setting, their use does not accurately represent the population poised to benefit the most from the intended applications of patient-specific hiPSC-derived vascular research. Neonates very rarely suffer from cardiovascular disease whereas more than 70% of the population over 60 do(16). With recent evidence indicating hiPSCs from older donors show increasing levels of methylation of genes associated with angiogenesis, it is unclear whether the research gathered from ND-hiPSC-EPs will translate to hiPSC-EPs sourced from and utilized by mature patients (MD-hiPSCs) (17).

To address this critical knowledge gap, we compared the functionality of hiPSC-EPs derived from neonate donors to those derived from donors over 30 years of age. We hypothesized that due to differences in methylation patterns correlated with advancing age, donor age would significantly influence the vasculogenic potential and cellular function of hiPSC-EPs. Although the methylation and gene expression trends reported by Lo Sardo et al. (17) were not preserved following differentiation into hiPSC-EPs, our data reveal clear functional differences. ND-hiPSC-EPs exhibited significantly greater vasculogenic potential, forming denser and more interconnected vascular networks in 3D environments compared to MD-hiPSC-EPs. We also observed other notable differences in differentiation efficiency, with MD-hiPSCs showing a higher EP yield, a decrease in cellular density, a lack of membrane-bound VE-Cadherin in 2D culture, and distinct methylation and gene expression profiles.

## Materials and Methods

### Cell Lines and Culture

We chose six induced pluripotent stem cell (hiPSC) lines to use in this study, 3 from neonate donors and 3 from mature donors. Neonate donors were under 1 week of age at the time of sampling, while mature donors were over 30 years old. The hiPSC lines from neonate donors included: iPS-DF19-9-11T.H, designated as neonate donor 1 (ND-1) (WiCell, Cellosaurus ID: CVCL_K054); WISCi007-C, designated as neonate donor 2 (ND-2) (WiCell, Cellosaurus ID: CVCL_VF54); and CBhiPSC6.2, designated as neonate donor 3 (ND-3) (Thermo Fisher, Cat# A18945, Cellosaurus ID: CVCL_RM92). The hiPSC lines from mature donors included: WTC11, designated as mature donor 1 (MD-1) (UCSFi001-A, Cellosaurus ID: CVCL_Y803); WTB, designated as mature donor 2 (MD-2) (Gladstone WTB hiPSC line); and EDi044-A, designated as mature donor 3 (MD-3) (Cellosaurus ID: CVCL_YB36). ND-1, ND-2, MD-1, and MD-2 were derived from skin fibroblasts, while ND-3 and MD-3 were derived from CD-34^+^ mononuclear blood cells.

All hiPSC lines were cultured on vitronectin-coated plates (Thermo Fisher) and maintained in Essential 8 (E8) medium (Thermo Fisher). Plates were coated with vitronectin according to the manufacturer’s protocol, using a 0.5 µg/mL solution prepared in Dulbecco’s phosphate-buffered saline (DPBS) and incubated at 37°C for 1 hour prior to hiPSC seeding. hiPSCs were colony passaged every 3–4 days at approximately 70–80% confluency using 0.5 mM Ethylenediaminetetraacetic acid (EDTA) in DPBS (Thermo Fisher). For routine culture, E8 medium was replaced daily. All cultures were maintained at 37°C in a humidified incubator with 5% CO₂.

### Endothelial Progenitor Differentiation Protocol

We adapted the differentiation protocol from Jalilian et al. to generate endothelial progenitors (EPs) from induced pluripotent stem cells (hiPSCs) (18,19). After culturing hiPSCs as described, we dissociated them into a single-cell suspension using Accutase (Innovative Cell Technologies) and plated them at a density of 10,000 cells/cm² on Geltrex-coated plates (Thermo Fisher) in Complete E8 medium supplemented with 10 µM Y-27632 (Selleck Chemicals).

One day after seeding, we replaced the media with Complete E8 without Y-27632. Two days after seeding, we replaced the media with differentiation-inducing media, consisting of Dulbecco’s Modified Eagle Medium Nutrient Mixture F-12 (DMEM-F12) (Cytiva) supplemented with N2 supplement (Thermo Fisher) and B27 without insulin (Thermo Fisher), along with 25 ng/mL Activin A (R&D Systems), 30 ng/mL Bone Morphogenic Protein 4 (BMP4, R&D Systems), and 0.15 µM 6-bromoindirubin-3-oxime (BIO, Selleck Chemicals). After an additional day, we replaced the differentiation media with fresh differentiation media supplemented with Activin A, BMP4, and BIO at the above concentrations.

Four days after seeding, we replaced the media with differentiation media supplemented with 2 µM SB431542 (Tocris) and 50 ng/mL Vascular Endothelial Growth Factor (VEGF, ACRO Biosystems). We continued replacing this media every day for two additional days with fresh differentiation media supplemented with VEGF and SB431541 at the above concentrations.

Seven days after seeding, CD-34⁺ hiPSC-EPs were isolated as follows. We incubated the differentiated cells with Accutase (STEMCELL Technologies) for 10 minutes at 37°C and dissociated them into single cells. After centrifuging at 300g for 5 minutes, we resuspended the resulting pellet in 200 µL of sorting buffer (2 mM EDTA, 0.5% bovine serum albumin in DPBS; Sigma-Aldrich) with 2 µL of CD-34-FITC antibody (Miltenyi Biotec). We incubated the cells at 4°C for 10 minutes, then filtered them three times using a 35 µm cell strainer (Corning) to remove cell and extracellular matrix aggregates.

To isolate CD-34⁺ hiPSC-EPs, we used fluorescence-activated cell sorting (FACS) (S3e; Bio-Rad). We determined CD34^+^ yield using population analysis tools from ProSort, the Bio-Rad software native to the S3e cell sorter, and FlowJo.

### Formulation of hiPSC-EP-laden hydrogels

We encapsulated hiPSC-EPs in collagen/norbornene-modified hyaluronic acid (NorHA) hydrogels using methods described previously by our group (19,20). We chose collagen NorHA hydrogels because of their tunable stiffness, cell affinity, and degradation properties allowing for optimized controlled vasculogenic conditions. Based on previous studies done by our group, we chose a cell density of 2.2 million cells/ml, a collagen density of 2.5 mg/ml, and a 25% crosslinking of the available norbornene groups between hyaluronic acid chains.

Shortly after sorting, we centrifuged the hiPSC-EPs at 300 g for 5 minutes and then resuspended them endothelial growth media 2 (EGM2; PromoCell Inc) with 10 µM Y-27632 and 1X Antibiotic Antimycotic (Sigma-Aldrich). We embedded the resuspended hiPSC-EPs in the Collagen/NorHA hydrogels as previously described (19,20)Briefly, we mixed 10X Medium 199 (ThermoFisher Scientific) with type 1 rat tail collagen dissolved in acetic acid (Corning) and neutralized the solution with 1 M sodium hydroxide (Sigma-Aldrich). We then added this solution to a mixture of NorHA, RGD (GCGYGRGDSPG) (2.06 mg/ml), and an enzymatically degradable peptide (KCGPQGIWGQCK) (2.21 mg/ml). Finally, lithium phenyl-2,4,6-trimethylbenzoylphosphinate (LAP) was incorporated into a final concentration of 0.025 wt% and served as the photo-initiator to crosslink the NorHA.

Next, we pipetted 50 μL of the neutralized collagen-NorHA-cell suspension into individual wells in an 18-well glass chamber slide (Ibidi) and allowed them to solidify for 30 minutes at 37°C and 5% CO₂. We then exposed the hydrogels to 365 nm light at 10 mW/cm² for 50 seconds (Omnicure Series 1500). The final hydrogels contained 1 wt% NorHA and 0.25 wt% collagen. Immediately after exposure to UV light, we covered the gels in 100 µL of EGM2 supplemented with 50 ng/mL VEGF, 10 µM Y-27632, and 1X Antibiotic Antimycotic. We replaced media daily with 100 µL EGM2 supplemented with 50 ng/mL VEGF for 7 days.

### Network Analysis

We visualized the vessel-like networks generated within the hydrogels by fixing the hydrogels in 4% paraformaldehyde (PFA) 7 days after formulation and staining with rhodamine-phalloidin (ThermoFisher Scientific). For network quantification, we acquired Z-stacks at 13 µm intervals on a spinning disk confocal microscope (Zeiss Axio Observer Z1 with Yokogawa CSU-X1M). At least 4 regions of interest across the full depth of the hydrogel were acquired per hydrogel sample, and at least three hydrogels were analyzed per experimental condition.

To analyze the total length, connectivity, and vessel diameter of the networks, we used a computational pipeline previously developed and described by our group (11). Briefly, we filtered and binarized confocal z-stacks of regions of interest of the hydrogel using ImageJ and then analyzed the stacks using a MATLAB script that converts the images into a nodal graph. This nodal graph is then used to determine the associated parameters of the capillary-like networks by analyzing the nodes (branch/endpoints) and links (vessels).

### 2D Cell Culture

hiPSC-EPs from all lines immediately after sorting were plated on vitronectin (0.5 µg/mL in DPBS) coated plates at 35,000 cells/cm^2^ and cultured in EGM2 with the initial seeding media supplemented with 10 µM Y-27632 and 1X Antibiotic Antimycotic. We replaced the media with fresh EGM2 daily for 7 days. After 7 days the cells were fixed using 4% PFA and stained with anti-VE-Cadherin at 1 µg/ml (Santa Cruz Biotechnology), anti-Calponin at 0.5 µg/ml (Abcam), and DAPI 1 µg/ml (R&D Systems) which are the recommended concentrations from the suppliers and imaged at 40x magnification. At least 4 regions of interest per well were imaged at random and at least 4 wells per cell line were imaged. The resulting cell counts and quantification were done using ImageJ.

### Epigenetic Analysis

We collected between 500,000 and 1,000,000 CD-34^+^ cells immediately after sorting and extracted genomic DNA using the PureLink™ Genomic DNA Mini Kit (Thermo Fisher) by following the manufacturer’s guidelines. Extracted and purified genomic DNA concentration was quantified using a Take3 and Cytation 3, then submitted to the University of Texas at Austin Genomic Sequencing and Analysis Core Facility (RRID:SCR_021713), which supplied us with the raw .idat files generated using the EPICv2 methylation assay (Illumina). All analyses were conducted in R Studio using standard workflows as described in the EPICv2 tutorial (21). IDAT files containing raw signal intensities were processed using the Sesame package in R for preprocessing and normalization (22). Initial quality control steps included assessing sample and probe-level metrics to remove low-quality or failed probes, background correction, and dye-bias normalization. ß values were extracted to quantify DNA methylation levels. Additional probe filtering was conducted to exclude SNP-overlapping and cross-reactive probes as well as to remove probes that were matched to sex chromosomes. Age prediction was performed using the predictAge function within Sesame, applying the Horvath skin and blood algorithm (23). Gestational age was predicted using the Knight algorithm and the methylclock package in R Studio (24,25). Discovery of differently methylated regions was done using the DMRcate package and recommended thresholds (26).

### RNA seq

Total RNA was collected from separate differentiations, purified with RNeasy Mini Kit (Qiagen) according to the manufacturer’s protocol, and submitted to the University of Texas Genomic Sequencing and Analysis Facility (Center for Biomedical Research Support, RRID: SCR 021713). Tagseq libraries were sequenced using the NovaSeq 6000 SR100 and preprocessed by the Bioinformatics Consulting Group at the University of Texas at Austin. Reads were then processed using the nf-core RNAseq pipeline (27). Briefly, reads were aligned to human reference genome GRCh38 using ‘STAR’, before quantifying gene counts using ‘Salmon’ (28,29).

Differentially expressed genes were determined using edgeR’s quasi-likelihood method (30). Differentially expressed genes were defined to have an adjusted p-value <0.05 (Benjamini-Hotchberg) and log-fold-change of ±1. Genes, ranked by log-fold-change, were used for gene-set enrichment analysis with fgsea package in R and the Gene Ontology Biological Processes, WikiPathways, KEGG Medicus, and Reactome gene sets (31–36). Significantly enriched pathways were defined as having adjusted p-values <0.05. Gene counts and associated code can be found at https://github.com/ZoldanLab/Aging.

### Statistical Analysis

Unless otherwise stated, comparisons between parameters were done using a one-way ANOVA with individual comparisons between ND-1 and MD-1, ND-2 and MD-2, and ND-3 and MD-3 with * indicating a p-value < 0.05, ** indicating a p-value < 0.001, *** indicating a p-value < 0.0001, and **** indicating a p-value < 0.00001.

## Results

### MD-hiPSCs Yield a Higher Percentage of CD34^+^ Cells After EP Differentiation

We chose 6 different human induced pluripotent stem cell (hiPSC) lines for use in this study, 3 neonate donor (ND-hiPSC) and 3 mature donor (MD-hiPSC) lines (**Fig. 1A**). Importantly, each ND-hiPSC line is matched to one MD-hiPSC line by both sex and somatic cell origin (ND-1 with MD-1, ND-2 with MD-2, and ND-3 with MD-3), with the number indicating the corresponding pair. We differentiated all 6 hiPSC lines into endothelial progenitors (hiPSC-EPs) with identical conditions and parameters following a previously described protocol (**Fig. 1B**) (19). All six hiPSC lines exhibited similar morphological progression throughout differentiation (**S.Fig. 1**). By day 5, all six lines expressed the early EP marker CD-34, which we used to isolate hiPSC-EPs through FACS.

**Figure 1.**
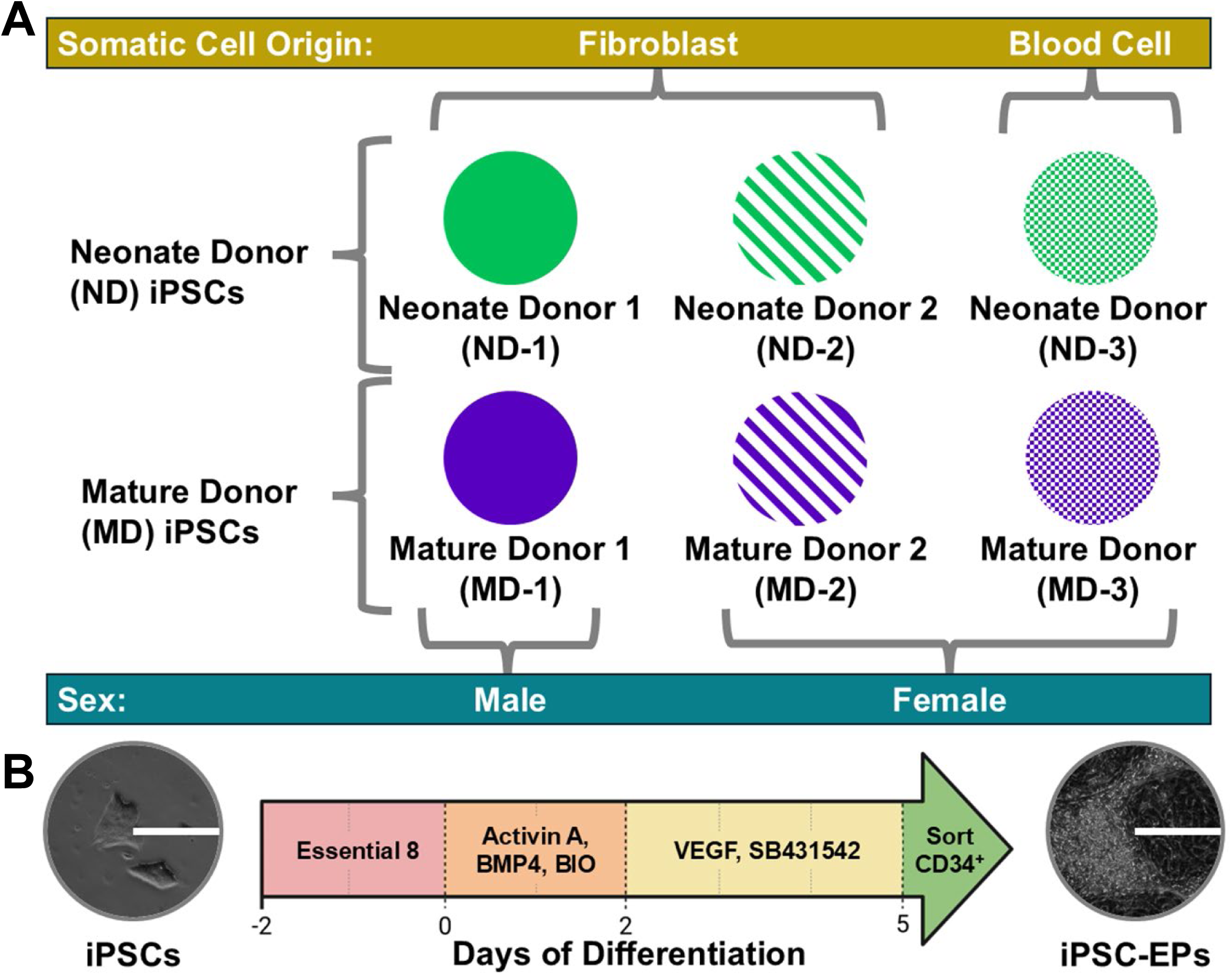
(A) Schematic illustrating the donor age, somatic cell source, and sex of the six iPSC lines used in this study. Colors and patterns assigned to each cell line are consistent across all figures**. (B)** Overview of iPSC differentiation to EPs. Representative bright field images of iPSCs colonies before mesoderm induction (left) and cells shortly before flow sorting (right). Scale bar: 200 µm.

Flow cytometry analysis revealed two distinct populations from each cell line (**Fig. 2A**). Since the positive populations were found in the same region across each cell line, we used identical gating to sort and quantify the CD-34^+^ cells. We found that MD-hiPSC lines consistently showed a significantly higher propensity for CD-34 expression (**Fig. 2B**). The average CD-34^+^ yield differentiated from MD-hiPSCs was significantly higher in both the matched pairs and higher than any ND-hiPSC line average. Differentiation yields of CD34^+^ cells for ND-hiPSCs ranged from 8.92-16.49% with a combined average of all replicates of 12.82 ± 5.32%, whereas the yields from all MD-hiPSCs ranged from 18.89-23.81% with an average of 20.68 ± 5.73%.

**Figure 2.**
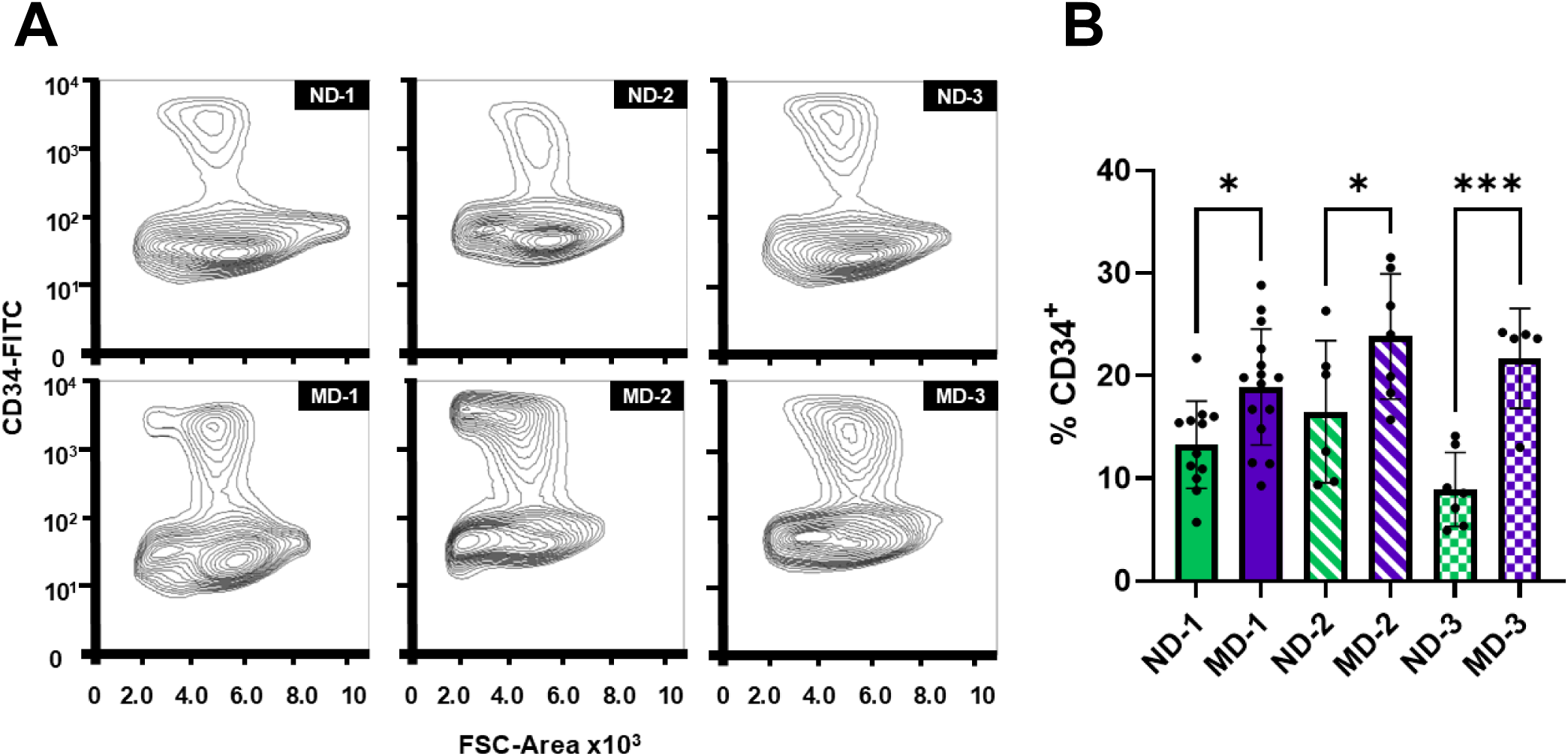
(A) Representative contour plots showing CD34⁺ and CD34⁻ populations from Day 5 differentiated iPSCs stained with anti-CD34. **(B)** Quantification of CD34⁺ cell yields across all six iPSC lines. MD-iPSC lines consistently produce a higher percentage of CD34⁺ cells compared to their matched ND-iPSC counterparts, as well as relative to all other ND-iPSC lines. One-way ANOVA with pairwise comparisons between ND-1/MD-1, ND-2/MD-2, and ND-3/MD-3. *p < 0.05, ***p < 0.0001. N = 5–15.

### Network Formation and Evaluation Shows Differences in ND- and MD-hiPSC-Eps

A key feature of hiPSC-EPs is their ability to form lumenized interconnected networks in 3D environments. To evaluate this capacity across the 6 chosen hiPSC-EP lines, we first encapsulated CD-34^+^ hiPSC-EPs in hydrogels composed of an interpenetrating network (IPN) of norbornene-functionalized hyaluronic acid (NorHA) and Collagen I. These hydrogels were formulated at cell densities and stiffness levels previously established by the Zoldan lab as the most optimal for vasculogenesis (19,20). After encapsulation and seven days of culture in EGM2 supplemented with VEGF, we quantified the resulting cellular structures to determine the vasculogenic potential using a computational pipeline developed by the Zoldan lab (**Fig. 3A-F**) (11). Interestingly, although the MD-hiPSC lines all showed a higher yield of EPs, (**Fig. 2A**) the EPs they produced all had reduced vasculogenic potential. Specifically, every ND-hiPSC-EP line developed a similar plexus architecture, whereas all MD-hiPSC-EPs failed to form a lumenized plexus (**Fig. 3G**).

**Figure 3.**
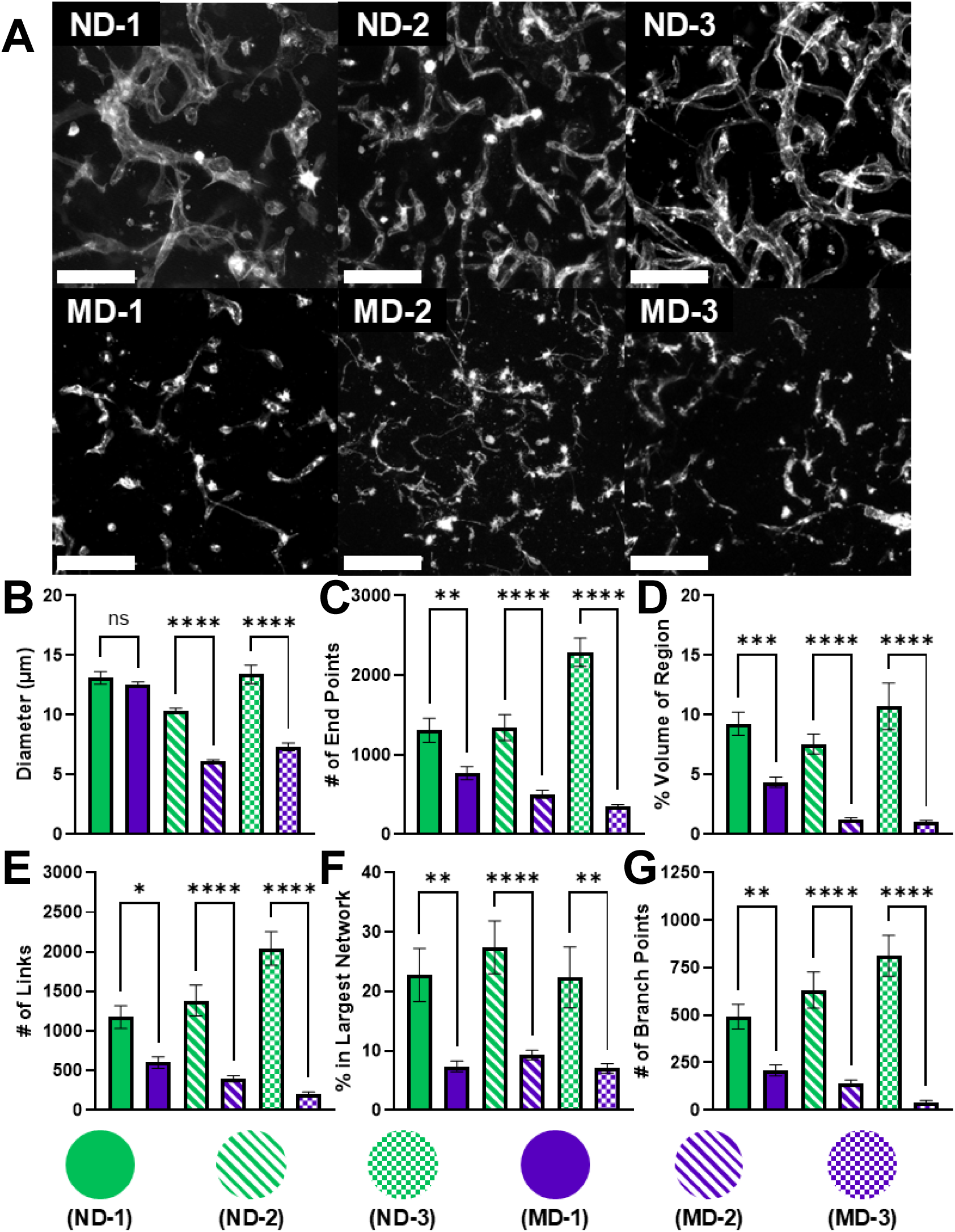
(A) Representative 50-100 µm deep Z-stack projection of the resulting micro vessel formed after encapsulating iPSC-EPs, differentiated from the six cell lines, in hydrogels and culturing for 7 days. Interconnected lumenized structures are observed in each ND-iPSC-EP line, while such structures are largely absent in the MD-iPSC-EP lines. **(B)** Average diameter of micro vessels (µm). **(C)** Average number of end points. These differences are reflected in the quantified vascular network metrics: **(D)** Total volume that is occupied by micro vessels (%). **(E)** The number of links between micro vessels. **(F)** The percent of micro vessels that are connected to the largest micro vessel. **(G)** The number of branch points within micro vessels. N=11-18. One-way-ANOVA with comparisons between ND-1/MD-1, ND-2/MD-2 and ND-3/MD-3. * = p<0.05, ** = p<0.01, *** = p<0.0001, **** = p<0.00001. Scale bar: 200 µm.

Unlike the ND-hiPSC-EPs, which generated extensive, lumenized, and highly interconnected vascular structures, the MD-hiPSC-EPs predominantly formed isolated clusters of cells, some with lumens but lacking interconnectivity (**Fig. 3D-G**). hiPSC-EPs sourced from MD-1 were the only MD-hiPSC-EPs with a similar average vessel diameter as ND-hiPSC-EPs and showed the most lumenized structures, however in all other metrics they underperformed compared to the ND-hiPSC-EPs (**Fig. 3A**). Between ND-2-hiPSC-EPs and MD-2-hiPSC-EPs (ND-2/MD-2) there is a 58% decrease in average vessel size and between ND-3-hiPSC-EPs and MD-3-hiPSC-EPs (ND-3/MD-3) there is a 45% decrease in average vessel size (**Fig. 3A**). Other metrics are similarly decreased in structures formed by MD-hiPSC-EPs with the number of end points decreasing by 41% between ND-1-hiPSC-EPs and MD-1-hiPSC-EPs (MD-1/ND-1), 83% (MD-2/ND-2) and 85% (MD-3/ND-3) reduction in the number of endpoints (**Fig. 3B**). Cellular structures only occupied 47% (MD-1/ND-1), 16.6% (MD-2/ND-2) and 9.2% (MD-3/ND-3) of the same volume (**Fig. 3C**). The number of links between vessels decreased by 49% (MD-1/ND-1), 89% (MD-2/ND-2) and 90% (MD-3/ND-3) (**Fig. 3D**). Interconnectivity was also severely impacted, with less than 10% of MD-hiPSC-EP formed vessels being connected to the largest vessel whereas ND-hiPSC-EPs were all between 22.4-22.9% connected (**Fig. 3 F)**. This lack of connectivity is further seen with a decrease of 57% (MD-1/ND-1), 93% (MD-2/ND-2), and 95% (MD-3/ND-3) in the number of branch points (**Fig. 3E**).

### 2D maturation of ND- and MD-hiPSC-EPs Results in Varied Morphologies

To investigate potential mechanisms underlying the reduced vasculogenic potential of MD-hiPSC-EPs, we examined the phenotypes of hiPSC-EPs in 2D culture. We previously demonstrated that CD34⁺ ND-hiPSC-EPs from ND-1 give rise to both endothelial cells (ECs), with clear membrane-bound VE-Cadherin, and smooth muscle cells (SMCs), expressing calponin, after 7 days in culture (19)

To determine whether MD-hiPSC-EPs undergo similar differentiation, we cultured EPs from both ND and MD lines in Endothelial Growth Medium 2 (EGM2) for 7 days and assessed cellular phenotype. All ND-hiPSC-EPs formed tight junctions with VE-Cadherin localized to the membrane, consistent with functional EC identity (**Fig. 4A**). In contrast, MD-hiPSC-EPs exhibited aberrant VE-Cadherin expression: MD-1-hiPSC-EPs and MD-2-hiPSC-EPs displayed weak signal restricted to the cytoplasm, while MD-3-hiPSC-EPs showed a mixed pattern of membrane-bound and cytoplasmic expression.

**Figure 4.**
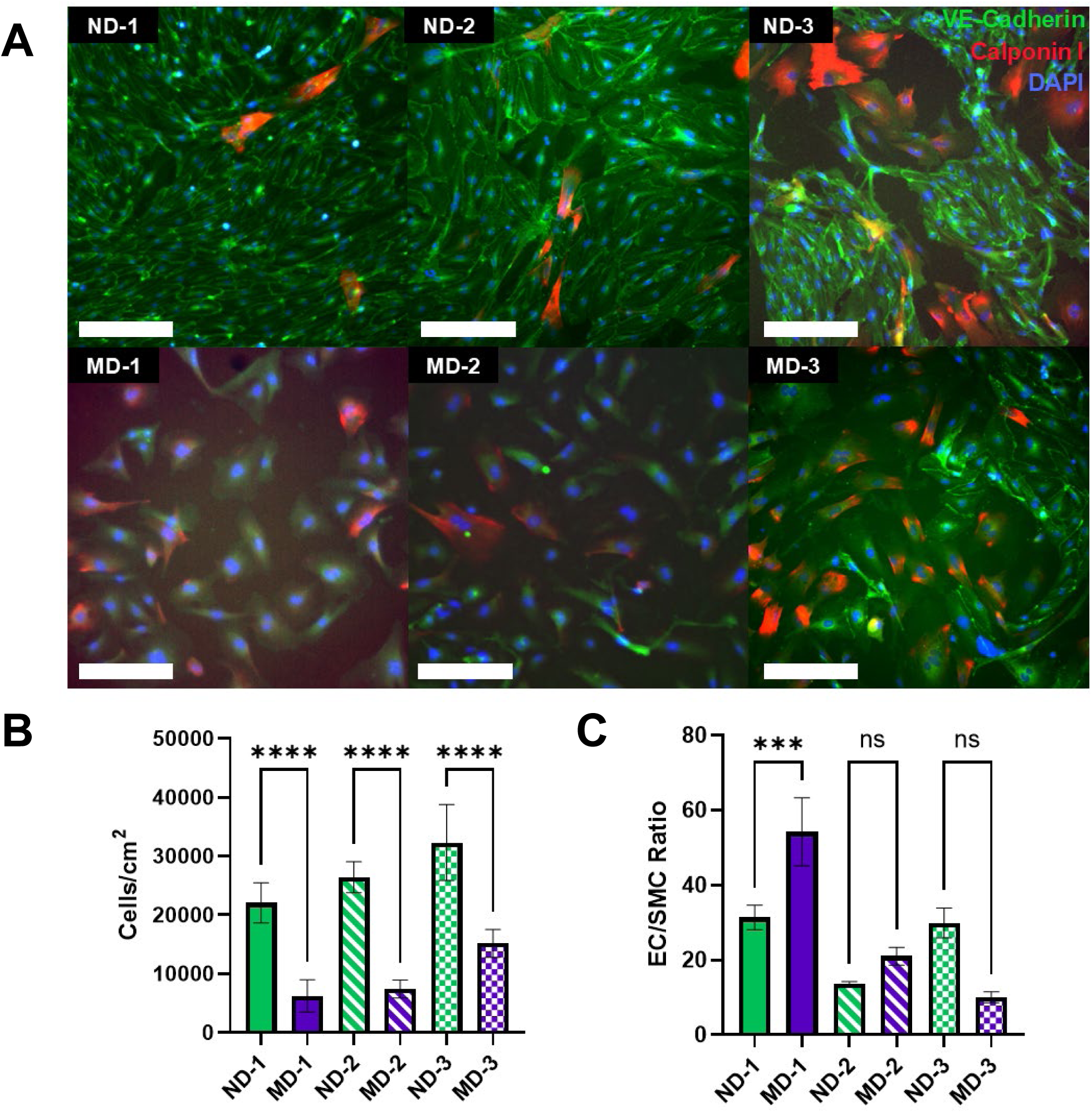
(A) Representative immunostaining images showing VE-Cadherin (green), nuclei (blue), and calponin (red) expression in endothelial cells differentiated from each of the six cell lines and cultured in 2D for 7 days in EGM2. Note the distinct membrane localization of VE-Cadherin and higher cell density in ND-iPSC-EPs, as well as the enlarged nuclei observed in MD-iPSC-EPs. **(B)** Quantification of cell density (cells/cm²). **(C)** Endothelial cell (EC) to smooth muscle cell (SMC) ratio, showing no significant differences. N=6-30. One-way-ANOVA with comparisons between ND-1/MD-1, ND-2/MD-2 and ND-3/MD-3. *** = p<0.0001, **** = p<0.00001. Scale bar: 250 µm.

Despite identical seeding densities, MD-hiPSC-EP cultures showed a statistically significant reduction in cell density compared to ND counterparts. However, the ratio of ECs to SMCs did not show a consistent trend between ND and MD lines (**Fig. 4B**).

### Methylation Signature Reveals Donor Background Differences

We next examined the epigenetic profiles of hiPSC-EPs derived from all six cell lines to determine whether there were consistent differences in DNA methylation patterns that persisted through differentiation. We examined the DNA methylation profiles at over 950,000 CpG sites using the EPICv2 methylation assay from Illumina. Methylation at each CpG site is binary, contributing a beta value of 1 if methylated and 0 if unmethylated. The average beta value then corresponds with the fraction of cells that are methylated at a specific CpG site in the tested population. Averaging beta values across all sites yields a global methylation level for each sample. Bulk methylation analysis of ß values revealed a clear trend: ND-1-hiPSC-EPs (0.673) and ND-3-hiPSC-EPs (0.657) showed statistically higher global methylation levels across all CpG sites compared to their respective MD counterparts (0.662 & 0.640) (**Fig. 5A&B**). In contrast, ND-2-hiPSC-EPs (0.647) exhibited lower global methylation than MD-2-hiPSC-EP (0.670); however, hiPSC-EPs differentiated from ND-2, ND-3, and MD-3 shared a similar methylation profile, and when compared to MD-3-hiPSC-EPs they showed a higher global methylation ß value.

**Figure 5.**
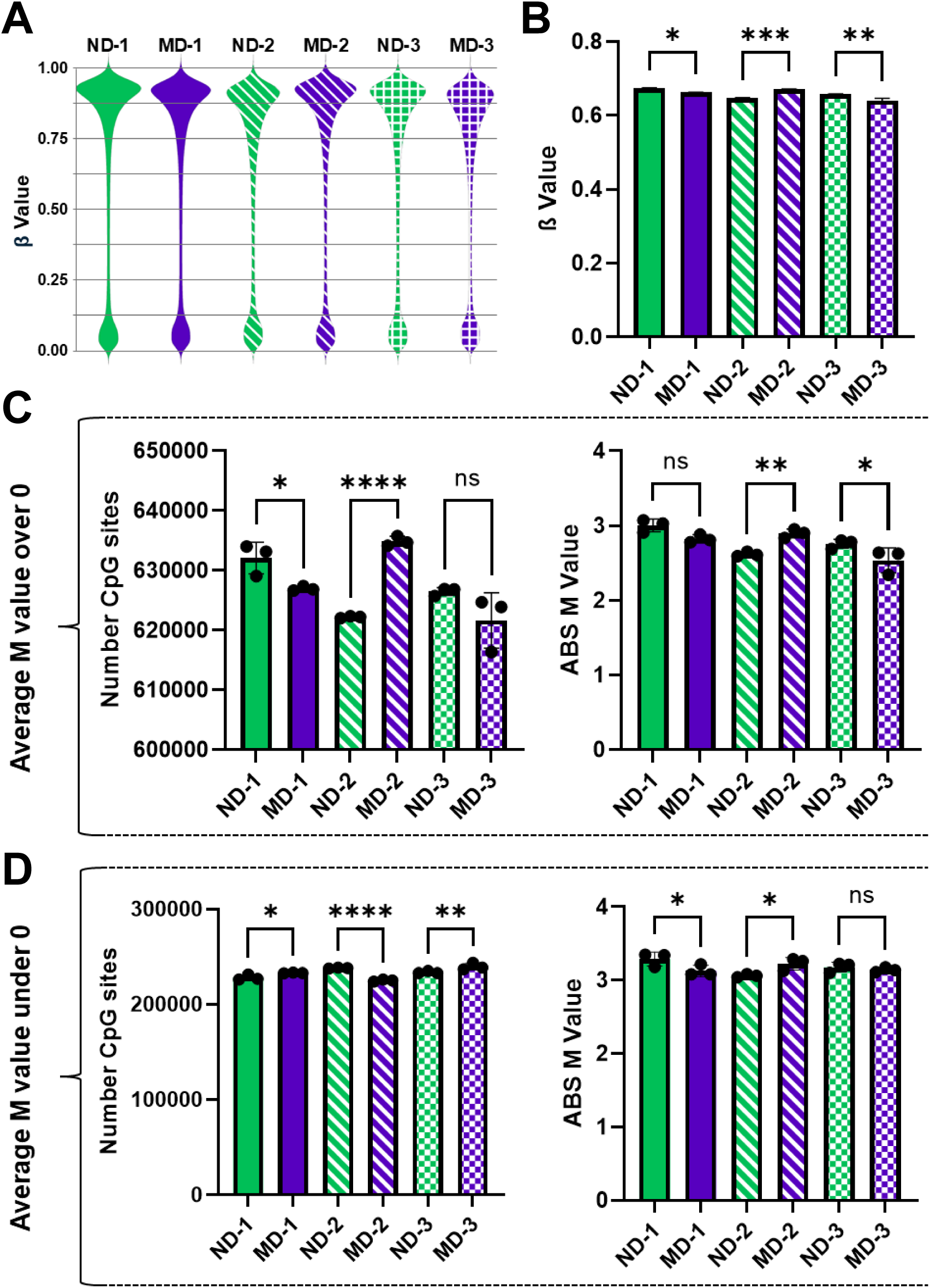
(A) Violin plots of beta values showing global methylation trends across all CpG sites, with ND1 and ND3 exhibiting higher overall methylation than their MD counterparts. **(B)** Total average ß methylation value for all CpG sites. **(C)** Total number of CpG sites and average absolute M value for all CpG sites with an M value over 0. **(D)** Total number of CpG sites and average absolute M value for all CpG sites with an M value below 0. N = 3. One-way-ANOVA with comparisons between ND-1/MD-1, ND-2/MD-2 and ND-3/MD-3. * = p<0.05, ** = p<0.01, *** = p<0.0001, **** = p<0.00001.

Converting ß-to-M values, which better capture variability in methylation states, we observed further differences in global methylation trends (37). An M-value of 0 for the CpG probe indicates an equal amount of methylated and unmethylated probe intensity. Filtering for only CpG sites with an M-value greater than 0, we found that ND-1-hiPSC-EPs and ND-3-hiPSC-EPs had a greater number of positive CpG sites (ND-1/MD-1 p-value = 0.045, ND-3/MD-3 p-value = 0.060) and a higher average positive M value (ND-1/MD-1 p-value = 0.08, ND-3/MD-3 p-value = 0.027) than their matched MD-hiPSC-EP lines (**Fig. 5C**). When filtered for M values below 0, both ND-1/MD-1 and ND-3/MD-3 comparisons revealed a statistically significantly reduction in the number of CpGs with ND-1-hiPSC-EPs also showing a significantly lower average M value (**Fig. 5D**). The lower M value magnitudes suggest a more homogenous unmethylated population. Notably, the ND-2-hiPSC-EP/MD-2-hiPSC-EP comparison deviated from this trend. Instead, the bulk methylation profile of ND-2-hiPSC-EPs more closely resembled that of ND-3-hiPSC-EPs and MD-3-hiPSC-EP, exhibiting an increased number of CpG sites with a positive M value relative to MD-3-hiPSC-EP. Overall, the global methylation profile was similar between ND-1-hiPSC-EPs, MD-1-hiPSC-EPs and MD-2-hiPSC-EPs, all derived from fibroblasts. Alternatively, ND-2-hiPSC-EPs, ND-3-hiPSC-EPs and MD-3-hiPSC-EPs grouped together, with ND-3 and MD-3 originating from blood-derived cells and ND-2, though reported as fibroblast-derived, aligning more closely with the blood-derived cells’ methylation signature.

This grouping observed in global methylation trends was further supported by multidimensional scaling (MDS) analysis of methylation variance between all cell lines (**S.Fig. 2A**). Dimension 1, representing the primary eigenvector and accounting for 34% of the total variance, showed strong grouping based on somatic cell origin, with the exception of ND-2-hiPSC-EPs, which deviated from this pattern and aligned more closely with ND-3-hiPSC-EPs and MD-3-hiPSC-EPs. Dimension 2, explaining 18% of the variance, revealed a weak separation between ND-hiPSC-EPs and MD-hiPSC-EP lines. Additional grouping among ND-hiPSC-EP, but not MD-hiPSC-EP, lines were observed in dimension 5 (11%) (**S.Fig. 2B**). No distinct grouping was identified in Dimension 3, which accounted for 15% of the variance. Grouping along higher-variance dimensions indicates that somatic origin is the primary contributor to methylation differences, while separation based on donor age background plays a secondary role in variation.

We further processed the CpG sites by mapping them to the hg38 reference genome and applied the DMRcate algorithm to identify differentially methylated regions (DMRs), which are clusters of CpGs with statistically significant differences in methylation. Each cell line was compared to every other cell line. A consistent trend emerged: comparisons between cell lines of similar somatic cell origin yielded the fewest DMRs. With the exception of ND-2/MD-2, all matched ND-MD pairs showed fewer DMRs than unmatched comparisons (**S.Fig. 3A-C**). Notably, the ND-2-MD-2 comparison yielded 3,279 DMRs—comparable but less than the ND-2-MD-1 comparison (6959 DMRs) and substantially more than the ND-2-MD-3 comparison (888 DMRs). However, the minimal DMR differences observed in the ND-1/MD-1 and ND-3/MD-3 comparisons support the strategy of using matched pairs to isolate methylation differences attributable to donor age. Given the closer similarity between ND-2-hiPSC-EPs and MD-3-hiPSC-EPs, we included both ND-2-hiPSC-EP/MD-2-hiPSC-EP and ND-2-hiPSC-EP/MD-3-hiPSC-EP (ND-2/MD-3) comparisons in subsequent analyses.

Typically, DMRs are assigned to nearby genes, and their average CpG ß value difference is reported alongside the regions identified. In our analysis, we identified several genes with associated DMRs that were common across all ND-MD comparisons (**S.Fig. 5D**). Among these, genes related to the WNT signaling pathway, such as *WNT3A* and *TRRAP*, exhibited less methylation in ND lines compared to MD lines (38,39). Notably, *GNAS*, a gene essential for early angiogenesis but not for vasculogenesis was found to be more methylated in ND lines compared to MD lines (40).

To refine our analysis, we expanded our search to include genes associated with DMRs shared across at least two of the ND-MD matched comparisons, provided the third comparison displayed a methylation difference of less than 0.1. This filtering approach identified 266 overlapping DMRs, several of which are associated with genes that are functionally relevant to endothelial cell biology and vasculogenesis (**Fig. 6A**). Shared trends emerged across these comparisons: ND-1/MD-1 and ND-3/MD-3 showed a consistent increase in methylation in ND lines, whereas ND-2/MD-2, ND-3/MD-3, and ND-2/MD-3 comparisons revealed overlapping sets of genes with decreased methylation in the ND lines (**Fig. 6A**).

**Figure 6.**
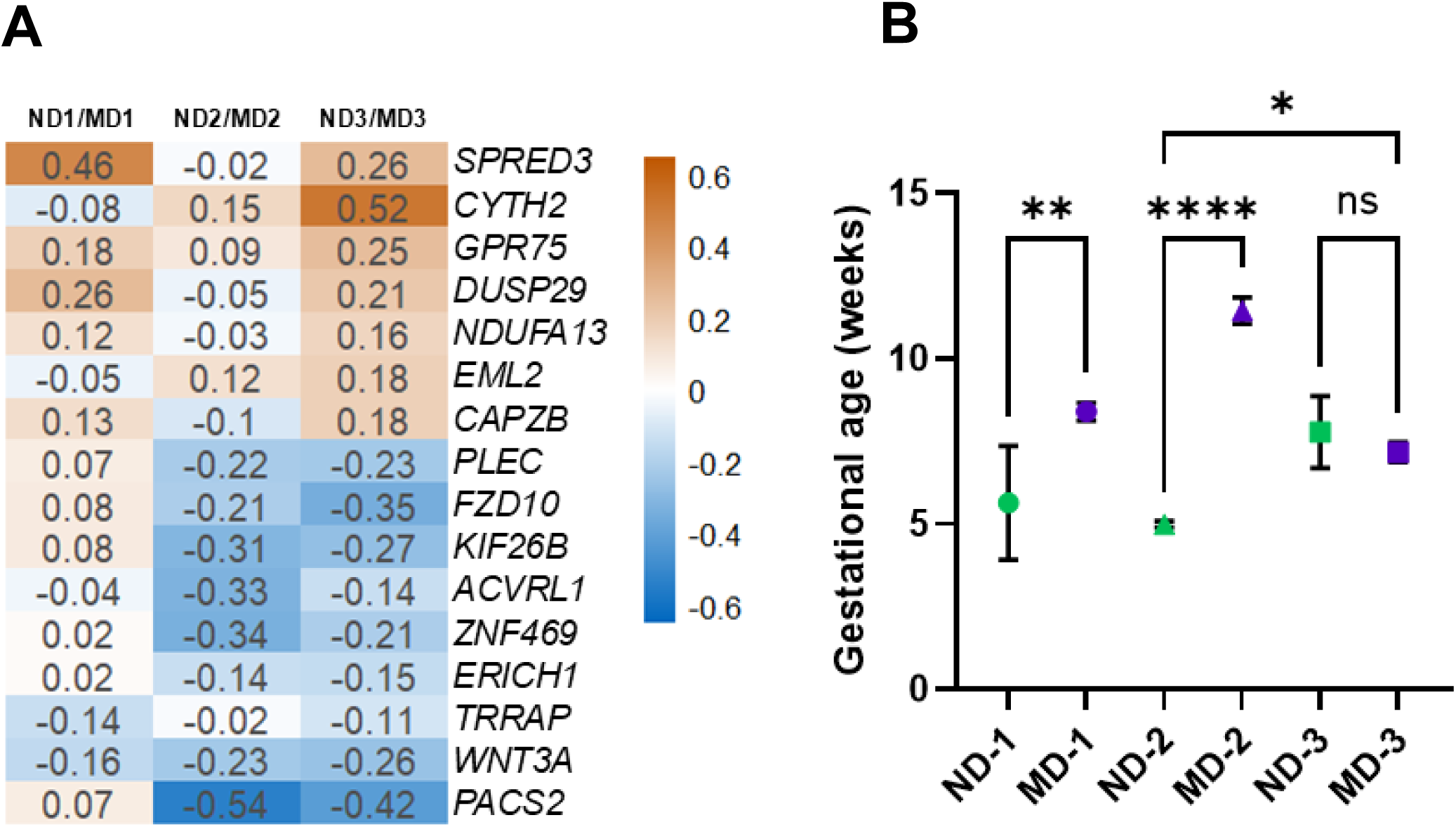
(A) Heatmap of relevant DMR-associated genes that are expressed in iPSC-EPs. Gene associated DMRs are shared across at least two ND/MD comparisons, with a methylation difference of less than 0.1 for the third ND/MD comparison. **(B)** Predicted gestational age based on the Knight et al. epigenetic clock, showing significantly lower predicted gestational age in ND-1/MD-1 and ND-2/MD-2 and significantly lower age in ND-2/MD-3. N = 3. One-way-ANOVA with comparisons between ND-1/MD-1, ND-2/MD-2, ND-3/MD-3 and ND-2/MD-3. * = p <0.05. ** = p 0.01, **** = p <0.00001.

Among the genes displaying shared differential methylation, several are known to play roles in processes critical for vascular development and function. These include *WNT3A*, *FZD-10* and *TRRAP* (WNT signaling) (38,39,41), *SPRED-3* (RAS/MAPK signaling) (42), *PLEC*, *CAPZB* and *KIF26B* (cytoskeletal organization and polarity) (43–45), *NDUFA13*, *TXNRD-2* and *PACS2* (mitochondrial function) (46–48), and *CYTH2 (ARNO)*, *ACVRL1*, *GPR75* (vascular development and endothelial signaling) (49–51) (**Fig. 5A**).

Lastly, we assessed the approximate epigenetic age of each line using the Horvath Skin and Blood clock which is a widely used model for predicting chronological age. All lines, except for ND-3 and MD-3, had a negative predicted age (**S.Fig. 4**), with no clear differences observed between cell lines. This suggested that although hiPSC-EPs, progressed through the differentiation process, similar to hiPSC, they retain fetal signatures. We therefore applied the Knight epigenetic clock to estimate gestational age. Using this model, ND-1-hiPSC-EPs (5.64 wks) and ND-2-hiPSC-EPs (5.00 wks) were predicted to be significantly younger than MD-1-hiPSC-EPs (8.40 wks) and MD-2-hiPSC-EPs (11.5 wks). ND-3-hiPSC-EPs (7.78 wks) and MD-3-hiPSC-EPs (7.17 wks) showed no statistical difference, although ND-2-hiPSC-EPs showed a significantly younger gestational age prediction than MD-3-hiPSC-EPs (**Fig. 5B**).

### Transcriptomic Profiling Reveals Somatic Cell, Age, and Sex Linked Expression Differences

We performed Tagseq RNA sequencing with subsequent normalization and analysis using EdgeR’s Trimmed Mean of M-values and quasi-likelihood generalized linear methods. Multidimensional scaling of resulting data was then used to evaluate the global variance between MD- and ND-derived EPs. In the MDS plot, dimension 1 accounting for 26% inter-sample variance showed separation of individual donors, with partial clustering of paired donors ND-1–MD-1-hiPSC-EPs and ND-3–MD-3-hiPSC-EPs, and an unexpected grouping of ND-2-hiPSC-EPs with MD-3-hiPSC-EPs, mirroring patterns previously observed in methylation-based clustering (**S.Fig. 5A**). Dimension 2 (18% of variance) broadly separated ND-hiPSC-EP and MD-hiPSC-EP groups, while dimension 4 (9%) showed grouping within the ND-hiPSC-EP samples only (**S.Fig. 5B**). No discernible clustering was observed in dimensions 3 (11%) or 5 (7%).

Hierarchical clustering of the most significantly differentially expressed genes (FDR < 0.05; (**S.Fig. 6A**) revealed separation by donor sex: ND-1-hiPSC-EPs and MD-1-hiPSC-EPs clustered together, while ND-2-hiPSC-EPs, ND-3-hiPSC-EPs, MD-2-hiPSC-EPs, and MD-3-hiPSC-EPs formed a separate branch. Among the most downregulated genes (**S.Fig. 6B**), clustering largely aligned with somatic cell origin except for ND-2-hiPSC-EPs. Clustering based on the most upregulated genes corresponded more closely with donor sex (**S.Fig. 6C**). Notable downregulated genes included *DDAH2*, involved in nitric oxide metabolism and vascular remodeling, and *TLN2*, a cytoskeletal adhesion protein, which were specifically downregulated in ND-1-hiPSC-EPs and MD-3-hiPSC-EP(52,53).

While broad trends showed separation by donor sex, pathway-level analysis using heatmaps of key gene sets largely confirmed separation by somatic cell origin, with the exception of ND-2-hiPSC-EPs, which consistently grouped with ND-3-hiPSC-EPs and MD-3-hiPSC-EPs. Notable exceptions to this trend included specific pathways “Regulation of Mitochondrial Gene Expression” (**Fig. 7A**), “Nitric Oxide Mediated Signal Transduction” (**Fig. 7B**), “Barbed End Actin Filament Capping” (**Fig. 7C**), and “Regulation of Extracellular Matrix Organization” (**Fig. 7D**). Of these, the only consistent separation of ND-hiPSC-EPs from MD-hiPSC-EPs was seen in “Regulation of Mitochondrial Gene Expression”. “Nitric Oxide Mediated Signal Transduction” was most active in ND-1-hiPSC-EPs and ND-2-hiPSC-EPs compared with all MD-hiPSC-EPs and ND-3-hiPSC-EPs, “Barbed End Actin Filament Capping” showed highest activity in ND-1-hiPSC-EPs and MD-1-hiPSC-EPs, while “Regulation of Extracellular Matrix Organization” was most enriched in ND-1-hiPSC-EPs and ND-2-hiPSC-EPs with all MD lines showing lower activation.

**Figure 7.**
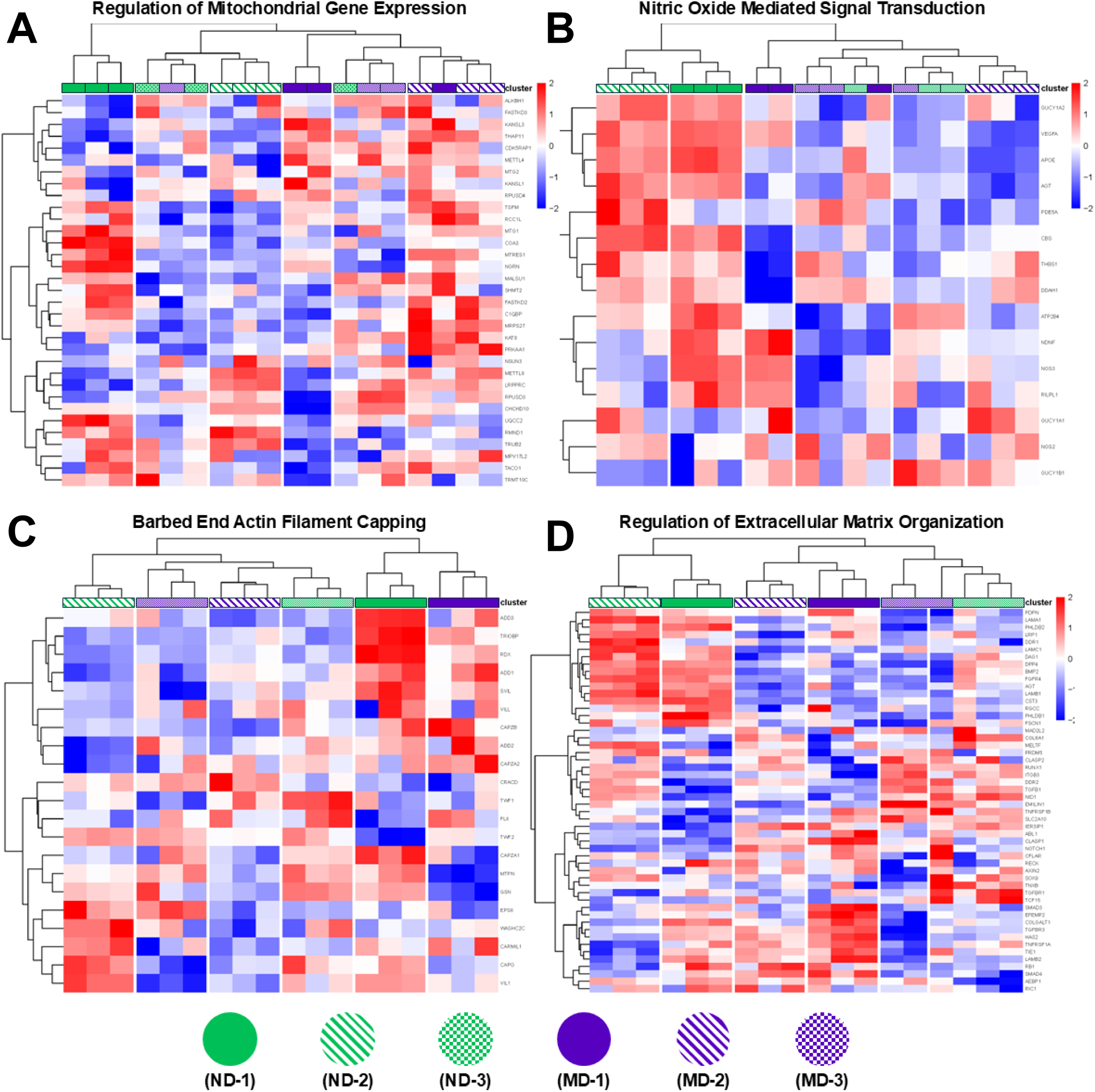
(A) Regulation of Mitochondrial Gene Expression shows consistent separation between ND and MD lines, with ND samples clustering separately. **(B)** Nitric Oxide Mediated Signal Transduction demonstrates elevated gene expression in ND1 and ND2 relative to other donors. **(C)** Barbed End Actin Filament Capping displays greater activation in ND1 and MD1 compared to other lines, suggesting donor-specific effects. **(D**) Regulation of Extracellular Matrix Organization reveals higher expression in ND1 and ND2, with ND3 clustering with MD lines but showing increased activation relative to other MD donors.

Additionally, although the pathway “VEGF and VEGFR Signaling Pathway” (**S.Fig. 7**) followed somatic cell origin grouping, it is worth noting the pronounced downregulation of several genes including, *ENG*, *GPC1*, *NCL*, and *SMARCA2*, in MD-1-hiPSC-EPs compared to ND-hiPSC-EPs, suggesting impaired extension and branching capacity which we observed *in vitro* (**Fig. 3**) (54–57).

Pairwise comparison between matched donors showed significant differential expression of *VEGFA*, *APOE*, and *AGT* between ND-2-hiPSC-EPz and MD-2-hiPSC-EPs, while *APOE* was significantly differentially expressed between ND-1-hiPSC-EPs and MD-1-hiPSC-EPs, suggesting nuanced differences in vascularization between donor pairs (**Fig. 7B**) (58–60).

Pathway enrichment of each ND and MD donor pair revealed significant upregulation of wound healing and cell-to-cell adhesion pathways in ND lines relative to MD lines. Additionally, pathways related to metabolism, hormone homeostasis, and signaling were also enriched in the ND--hiPSC-EP (**S. Fig. 8-10**).

## Discussion

Many hiPSC lines used in cardiovascular research originate from neonate donors, but future autologous applications will primarily rely on adult donor cells. This disparity warrants attention because prior studies show donor-age-associated differences in hiPSCs. With increasing donor age comes an increase in genomic and mitochondrial mutations as well as methylation and gene expression variability (17,61). While these differences do not appear to impair pluripotency or differentiation capacity, their influence on the function of fully differentiated progeny remains unclear (9). To ensure the translational relevance of past research, it is critical to determine whether hiPSC-derived cells from neonatal and adult donors differ in functionality.

Our study addresses this knowledge gap by directly comparing the vasculogenic potential of hiPSC-derived endothelial progenitors (hiPSC-EPs) derived from neonatal donors (ND) and mature donors (MD) in both 2D and 3D culture. Notably, we observed marked phenotypical differences in both 2D and 3D culture conditions (**Fig. 3&4**). All MD-hiPSC-EPs failed to generate dense, interconnected vascular networks in 3D culture, possibly through distinct mechanisms. We further characterized transcriptomic and epigenetic profiling and found inter- and intra-group differences, with somatic cell origin potentially driving some of the observed variance. However, consistent differences between ND and MD-hiPSC were also evident. Together our observations of the vasculogenic deficiency in MD-hiPSC-EPs and differences in methylation profiles and gene expression suggest that while MD-hiPSCs differentiate into EPs, MD-hiPSC-EPs may be functionally impaired through several candidate mechanisms.

We first considered whether the observed functional deficits could be attributed to contamination by CD34-expressing non-endothelial progenitor lineages. However, we observed high expression of canonical EP markers, including CD34 from FACS, and *CD31*, and *CDH5* from RNA sequencing, alongside low or undetectable expression of alternative CD34⁺ lineage markers (62). This gene expression profile indicates that the differentiated populations were indeed composed of EPs.

Since not all EPs are functionally or transcriptionally equivalent (19), we next assessed the functional and transcriptional variance among EPs derived from each of the six selected hiPSC lines. Our first indication that such differences may exist was consistently higher CD34⁺ cell yields from MD-hiPSCs (**Fig. 2**). These data suggest intrinsic differences in lineage specification or expansion capacity.

To further investigate the basis of these differences and characterize the heterogeneity within the EP populations, we examined the DNA methylation profiles. Except for ND-2/MD-2, ND-hiPSC-EPs consistently exhibited higher average methylated values (**Fig. 5**) compared to their matched MD-hiPSC-EP counterparts. However, this trend was only observed within matched cell line pairs and does not, on its own, explain the observed functional differences between MD- and ND-hiPSC-EPs. We thought there may be an association between the decrease in methylation in the MD lines, to hypomethylation that occurs during hiPSC differentiation progression (63). This connection would suggest that differences in methylation may reflect underlying disparities in differentiation progression, with MD-hiPSC-EPs potentially undergoing accelerated epigenetic remodeling.

Since hiPSC differentiation recapitulates aspects of gestational development, (64) we examined whether predicted gestational age differed between ND- and MD-hiPSC-EPs. Using the Knight algorithm (25), which predicts gestational age based on methylation levels at 148 CpG sites, we found that MD-hiPSC-EPs exhibited more developmentally advanced epigenetic signatures (**Fig. 6B**). The predicted age ranged from 5-7 weeks in ND-hiPSC-EPs and 7-12 weeks in MD-hiPSC coinciding with the developmental stage of vasculogenic to angiogenic transitioning in early human development (65–67). These findings suggest that the more advanced predicted gestational age of MD-hiPSC-EPs may reflect a shift away from the vasculogenic developmental stage beyond and instead toward a more angiogenic state.

Supporting this hypothesis, the cell line with the most advanced predicted gestational age, MD-2, developed cellular networks in 3D cultures that closely resembled those generated by human umbilical vein endothelial cells (HUVECs) in 3D culture . HUVECS are developmentally more mature (68) than hiPSC-EPs and show extensive branching but limited lumen formation in NorHA-based hydrogels similar to the ones used in this study.

To further validate these developmental trends and identify gene-specific contributors, we next analyzed differentially methylated regions (DMRs) that varied between matched pairs as well as the total mRNA expression in EPs derived from the 6 cell lines. Consistently across all three matched pairs, we observed increased methylation in MD-iPSC-EPs of WNT signaling pathway genes which are critical for early mesoderm induction (**Fig. 5A**). This epigenetic repression of WNT signaling genes suggests that MD-hiPSC-EPs may be developmentally more advanced and further removed from the initial mesodermal priming stage than their ND-hiPSC-EPs counterparts.

Pathway enrichment analysis based on methylation variance and RNA expression frequently clustered cell lines according to their somatic cell origin, with the exception of ND-2 which grouped with the blood-derived hiPSC-EPs ND-3 and MD-3 (**S.Fig. 2,5,6**). However, paired comparisons in both DMRs and gene expression revealed molecular signatures that aligned with phenotypic differences observed in 3D culture. MD-1-hiPSC-EPs formed multicellular clusters with occasional lumen-like structures but failed to interconnect to the extent observed in all ND-hiPSC-EPs. This phenotype may be linked to reduced VEGF/VEGFR signaling pathway genes (**S.Fig 7**), along with increased actin filament capping, as suggested by both methylation and transcriptomic data (**Fig. 6&7**). Notably, although ND-1-hiPSC-EP clustered with MD-1-hiPSC-EP in these pathway analyses, direct comparisons showed that ND-1-hiPSC-EP exhibited a higher expression of VEGF-related genes, along with reduced actin capping activity. In addition, ND-1-hiPSC-EP did not cluster with MD-1-hiPSC-EP in pathways related to ECM organization and nitric oxide (NO)-mediated signal transduction (**Fig. 7**). NO deficiency is often associated with endothelial aging, and may contribute to the decreased vessel formation capacity observed in MD-1-hiPSC-EP (69).

MD-2-hiPSC-EPs exhibited extensive cellular protrusions but failed to form any lumenized structures in 3D culture. Although the exact mechanism remains undetermined, this phenotype may result from reduced cytoskeletal organization, as suggested by methylation data (**Fig. 6**), or from altered regulation of the extracellular matrix, as indicated by both RNA-seq and epigenetic profiles (**Fig. 6&7**).

MD-3-hiPSC-EPs exhibited a hybrid phenotype, displaying characteristics observed in both MD-1-hiPSC-EPs and MD-2-hiPSC-EPs, with significantly less lumen-like structures compared to MD-1-hiPSC-EPs but with more branching and extension (**Fig. 4**). However, MD-3-hiPSC-EPs still failed to form interconnected lumenized structures to a degree comparable to those generated by ND-hiPSC-EPs. Like MD-1-hiPSC-EPs, MD-3-hiPSC-EPs showed reduced enrichment in NO-mediated signaling and ECM organization pathways. Furthermore, MD-3-hiPSC-EPs exhibited higher methylation of several genes associated with capillary morphogenesis (**Fig. 6A**). These differences may underline their overall limited ability to form interconnected and lumenized structures.

Transcriptomic differences that clearly distinguished ND from MD-hiPSC-EPs, identified differential regulation in pathways associated with mitochondrial gene expression. Specifically, MD-hiPSC-EPs exhibited higher overall activation of mitochondrial regulation gene expression compared to ND-hiPSC-EPs (**Fig. 7**). This differential expression suggests a potential divergence in mitochondrial function. Prior studies have shown that mitochondrial DNA mutations increase with age and persist through hiPSC reprogramming (61). These mutations contribute to the heterogeneity of hiPSC lines and contribute to functional variation in the mitochondria of hiPSC-derived cells (70). Given that hiPSCs and EPs rely primarily on glycolysis during early stages, such differences might not be readily apparent until later stages requiring oxidative phosphorylation. Both glycolysis and oxidative phosphorylation are essential for lumen extension, and impaired or varied mitochondrial activity in MD hiPSC-EPs may partially explain their consistently reduced vasculogenic capacity (71).

We did not observe patterns consistent with those reported by Lo Sardo et al., who found that donor age correlated with persistent increases in methylation of *TEAD3*, *ADGRL4*, and *ZNF217* (17). In our study, we observed a slight decrease in the average methylation of a DMR associated with *TEAD3* in ND-2-hiPSC-EPs compared to MD-2-hiPSC-EPs, but no differences were observed in the other matched pairs. For *ADGRL4*, methylation was higher in MD-1-hiPSC-EPs relative to ND-1-hiPSC-EPs, with no significant differences in the other pairs. We did not detect any donor-age specific differences in methylation around *ZNF217*. These discrepancies may reflect differences in experimental context: while Lo Sardo et al. examined the epigenetic landscape of undifferentiated hiPSCs from adult donors aged 21 years and older, our study utilized differentiated hiPSCs and compared neonate with mature donors.

Taken together, our findings suggest that donor variability, including donor age, must be carefully considered in the development of hiPSC-EP-based vascular therapies. While both ND- and MD-derived hiPSC-EPs can undergo EC and SMC differentiation, MD-derived cells more frequently exhibit impaired formation of well-connected, lumenized vascular networks in vitro. These functional deficits may stem from differences in mitochondrial activity, epigenetic regulation, or developmental priming. Our data indicate that hiPSC-EPs derived from adult donors may be less capable of forming *de novo* vasculature, potentially limiting their utility in regenerative applications.

It remains to be determined whether the vasculogenic deficits observed in MD-derived hiPSC-EPs can be rescued, whether through co-culture with supportive cell types, supplementation with additional growth factors, or via paracrine signaling *in vivo*. Despite their limited ability to form interconnected vascular networks *in vitro*, MD-hiPSC-EPs shared many molecular and functional features with ND-hiPSC-EPs. This raises the possibility that MD-iPSC-EPs may still be capable of integrating with host vasculature or contributing to models mimicking angiogenesis but not vasculogenesis. Collectively, these findings highlight the complexity hiPSC donor background introduces, particularly following differentiation, and underscore the need to carefully consider donor-related variables when designing translational strategies for hiPSC-based tissue engineering.

## Acknowledgements

We would like to thank Dr. Adrianne Rosales and Katie Halwachs for their support with the NorHA Collagen hydrogel material formulation as well as Brett Stern for optimizing the hiPSC-EP differentiation method.

---

**Supplemental Figure 1.**
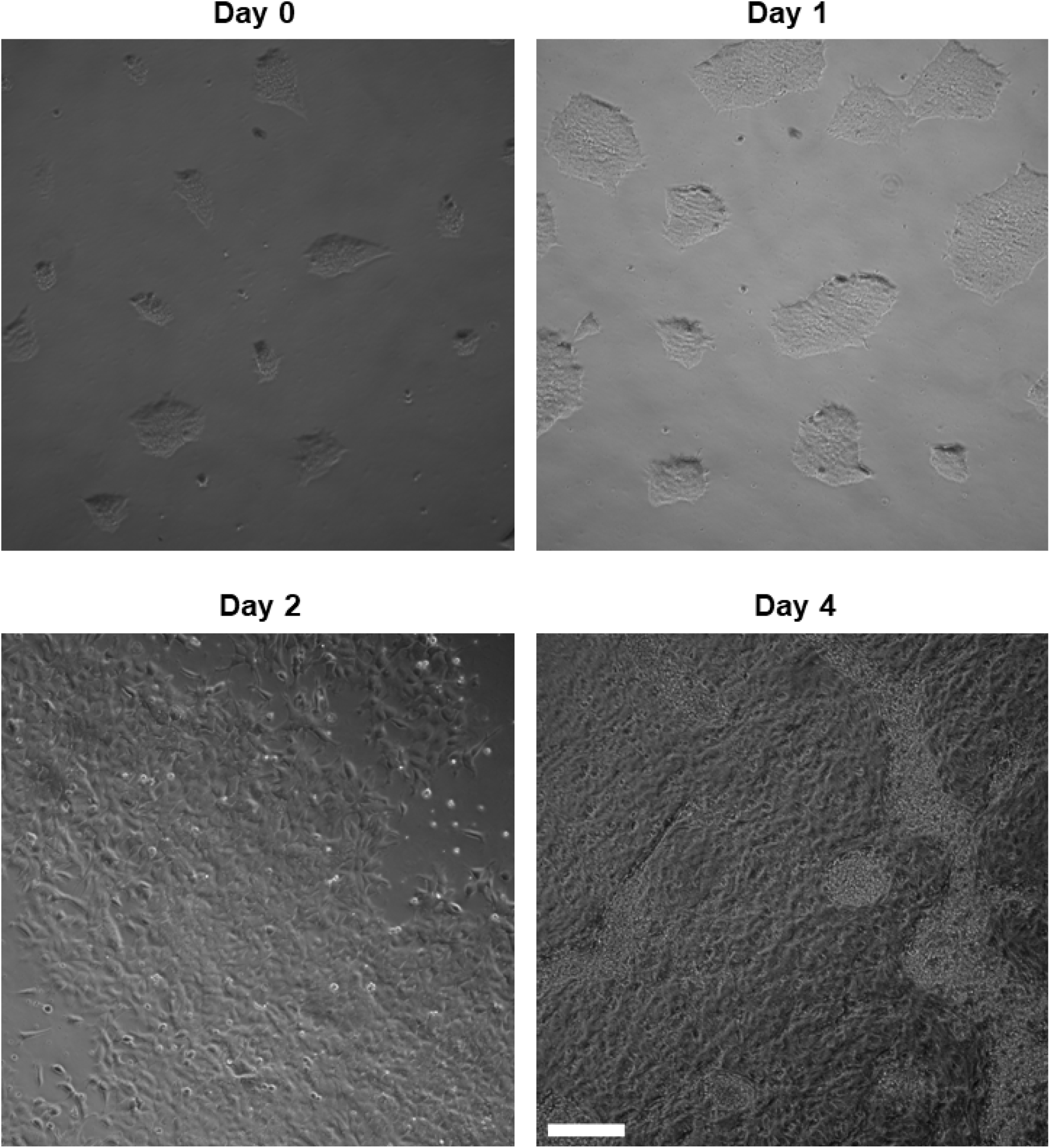
Representative images of iPSC to EP differentiation with formation of a mesodermal monolayer and enrichment of endothelial progenitor population. Scale Bar is 200 µm.

**Supplemental Figure 2.**
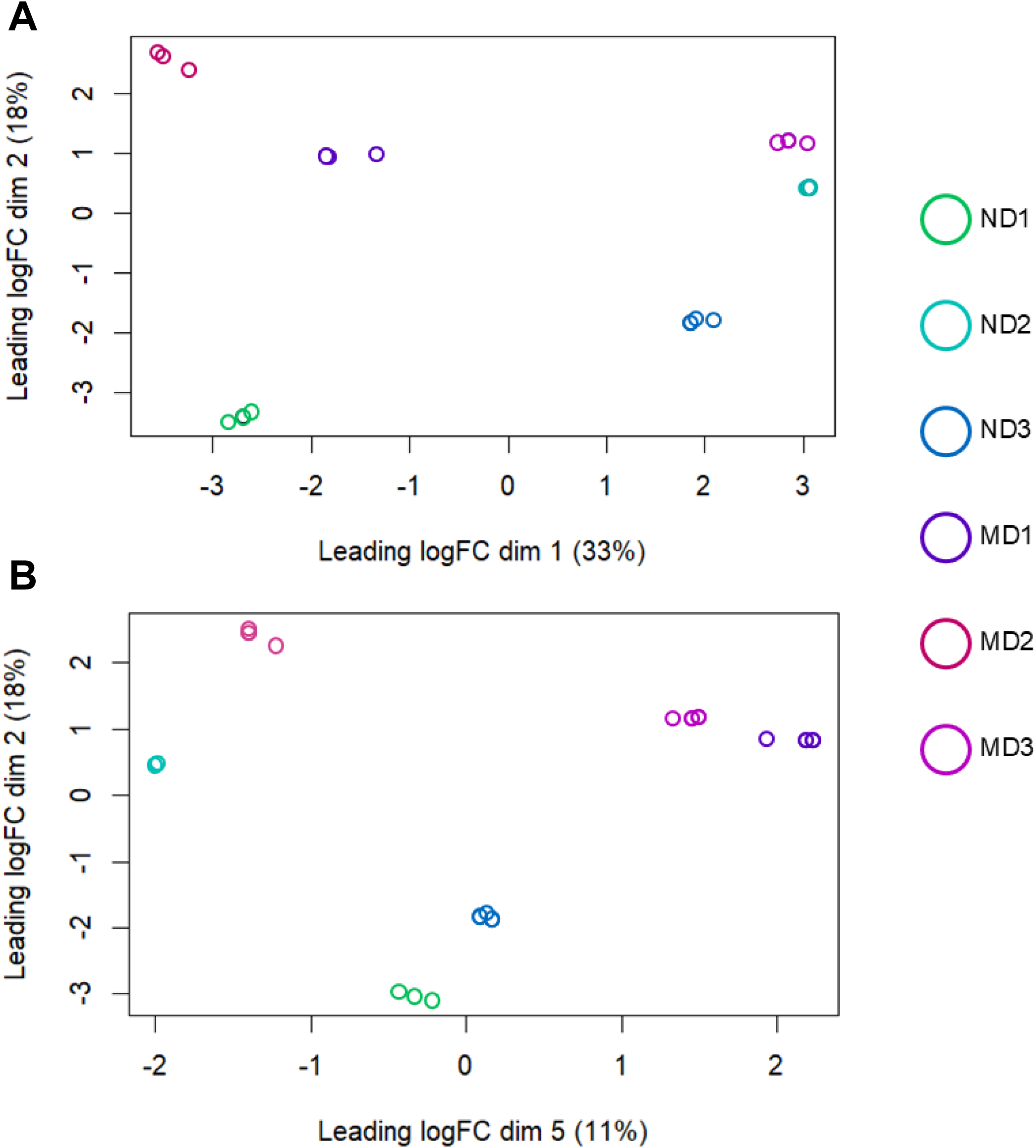
(A) Multidimensional scaling (MDS) plot of dimensions 1 and 2. Dimension 1 (34% variance) shows strong grouping by somatic cell origin, while dimension 2 (18%) reveals weaker separation by donor age. **(D)** Dimensions 2 and 5 (11%) showing ND grouping but not MD in dimension 5.

**Supplemental Figure 3.**
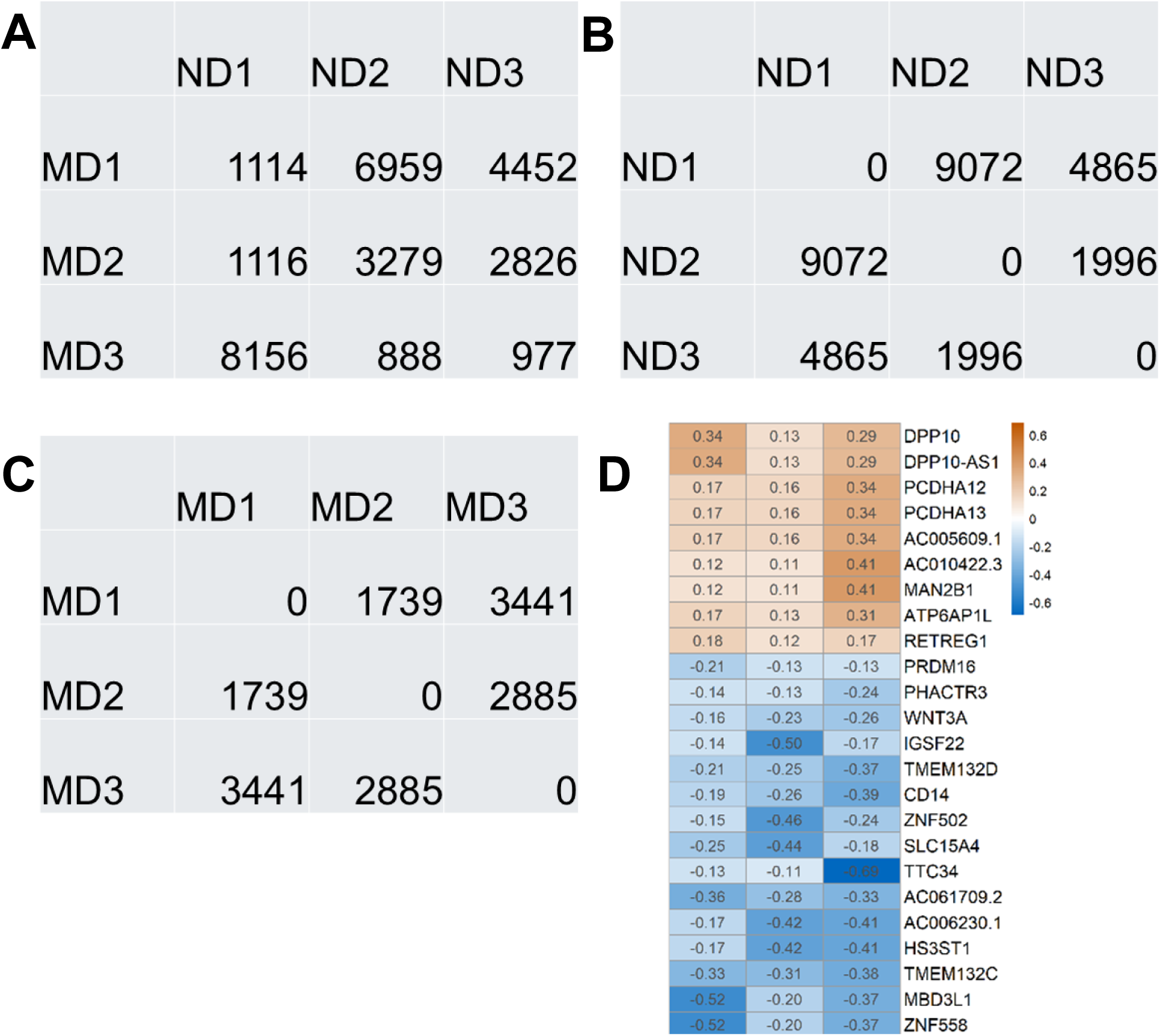
(A) Number of differentially methylated regions (DMRs) between each neonate and mature cell line. **(B)** Number of DMRs between each neonate line and every other neonate line. **(C)** Number of DMRs between each mature line and every other neonate line. **(D)** Heatmap of DMR-associated genes common to all ND/MD comparisons with an average CpG value difference greater than 0.1.

**Supplemental Figure 4.**
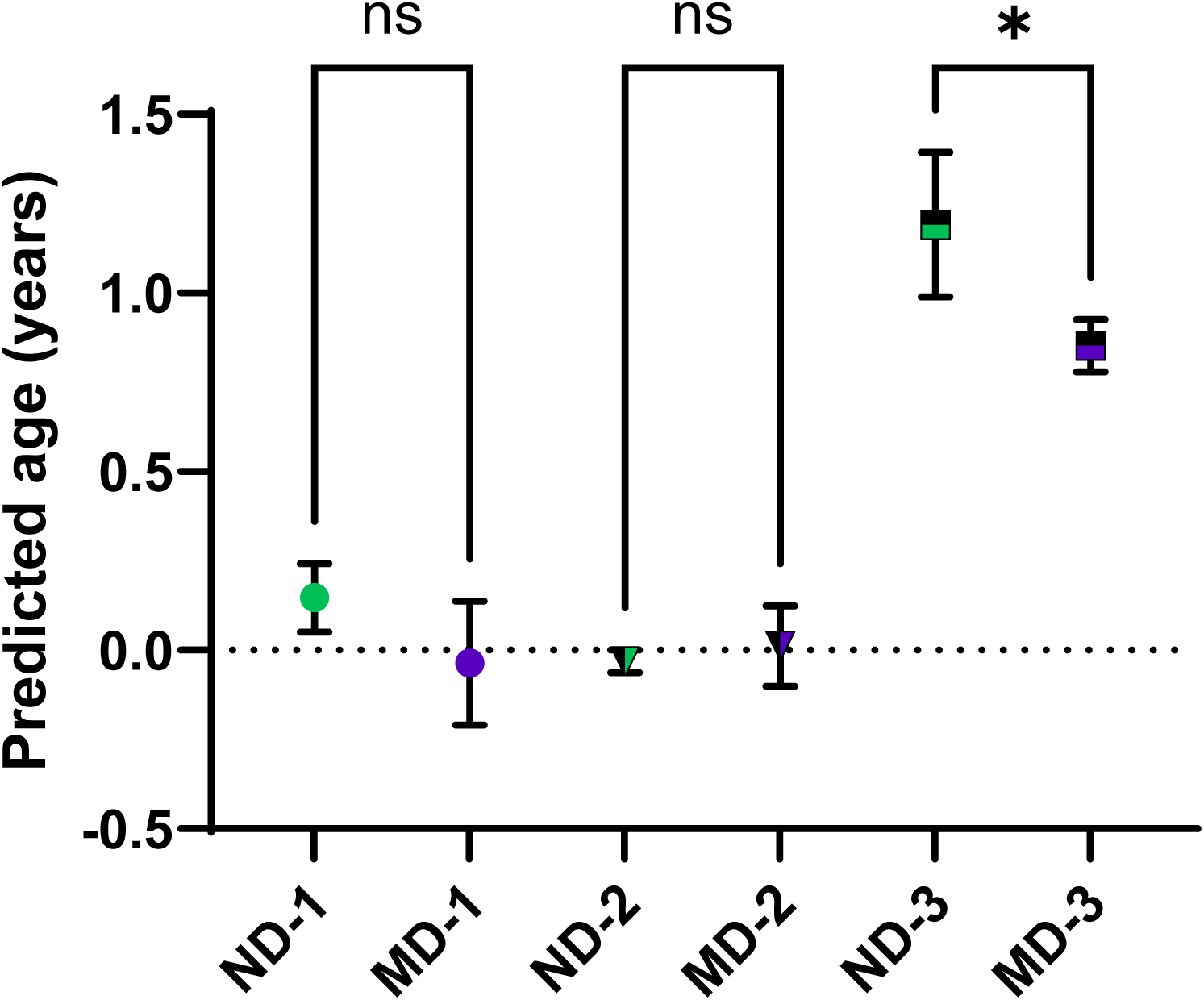
(A) Horvath predicted age shows no clear trends, with some lines appearing to have a negative predicted age similar to undifferentiated iPSCs. N = 3, * = p <0.05.

**Supplemental Figure 5.**
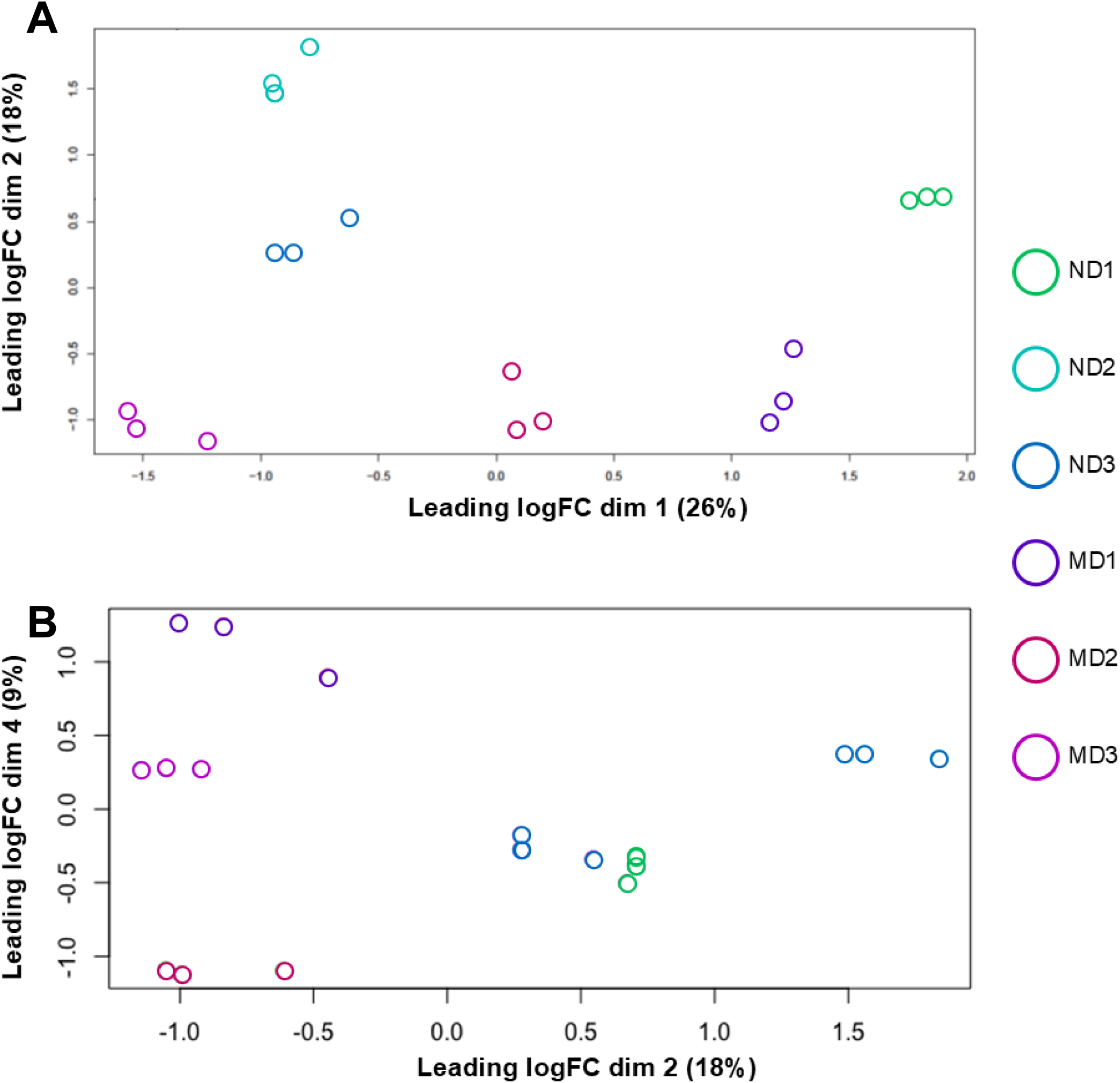
(A) MDS plot of dimensions 1 and 2 showing strong grouping along somatic cell origins in dimension 1 and weak grouping among age groups in dimension 2. **(B)** MDS plots of dimension 2 and 4 showing strong grouping in dimension 2 and ND grouping in dimension 4 but not MD.

**Supplemental Figure 6.**
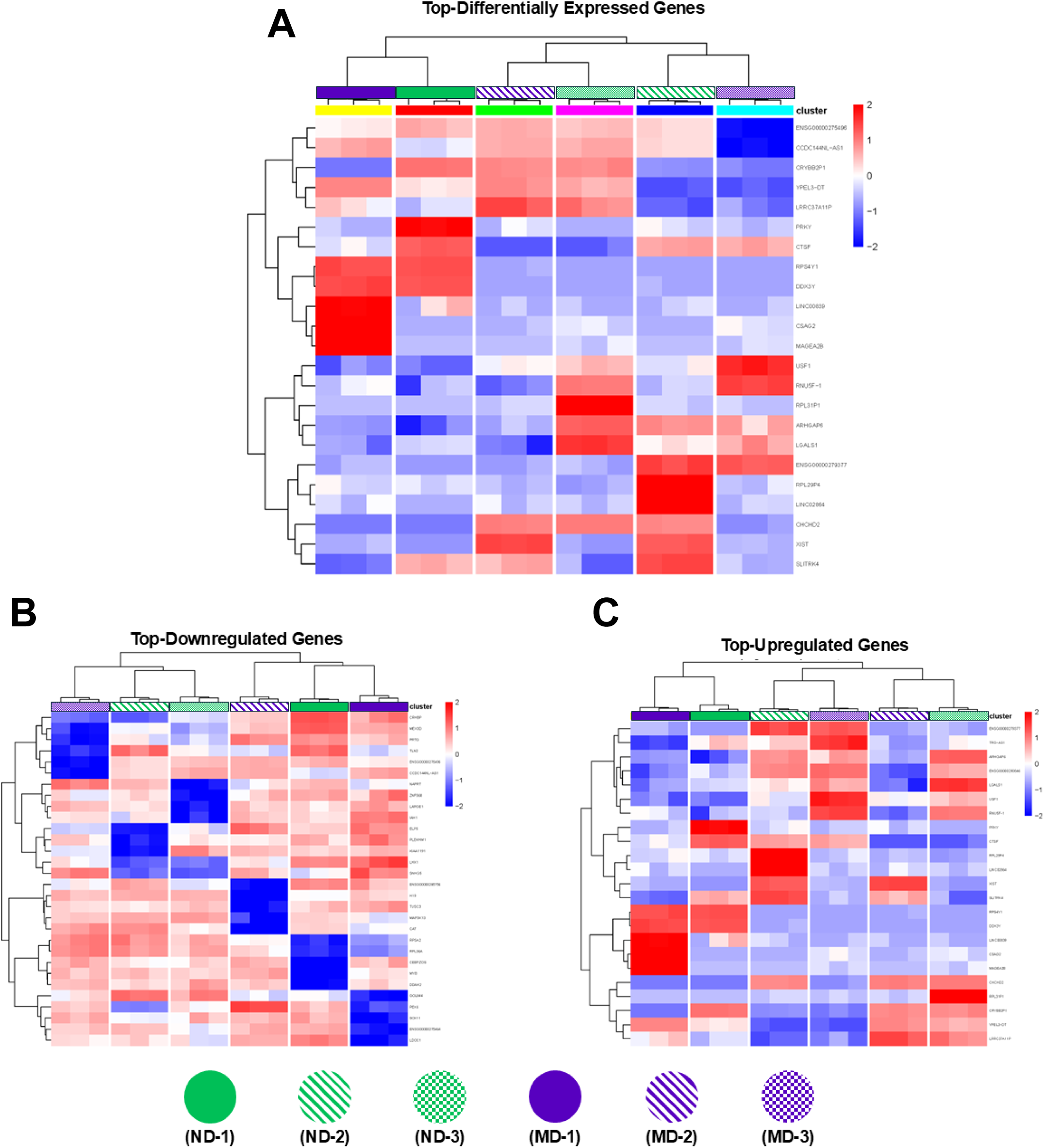
(A) Bulk-RNA Tagseq expression differences between Neonatal and Mature Donors. Clusterchart of top (p-value <0.05) log-transformed counts-per-million genes. **(B)** Clusterchart of top downregulated (p-value <0.05) log-transformed counts-per-million genes. **(C)** Clusterchart of top upregulated (p-value <0.05) log-transformed counts-per-million genes.

**Supplemental Figure 7.**
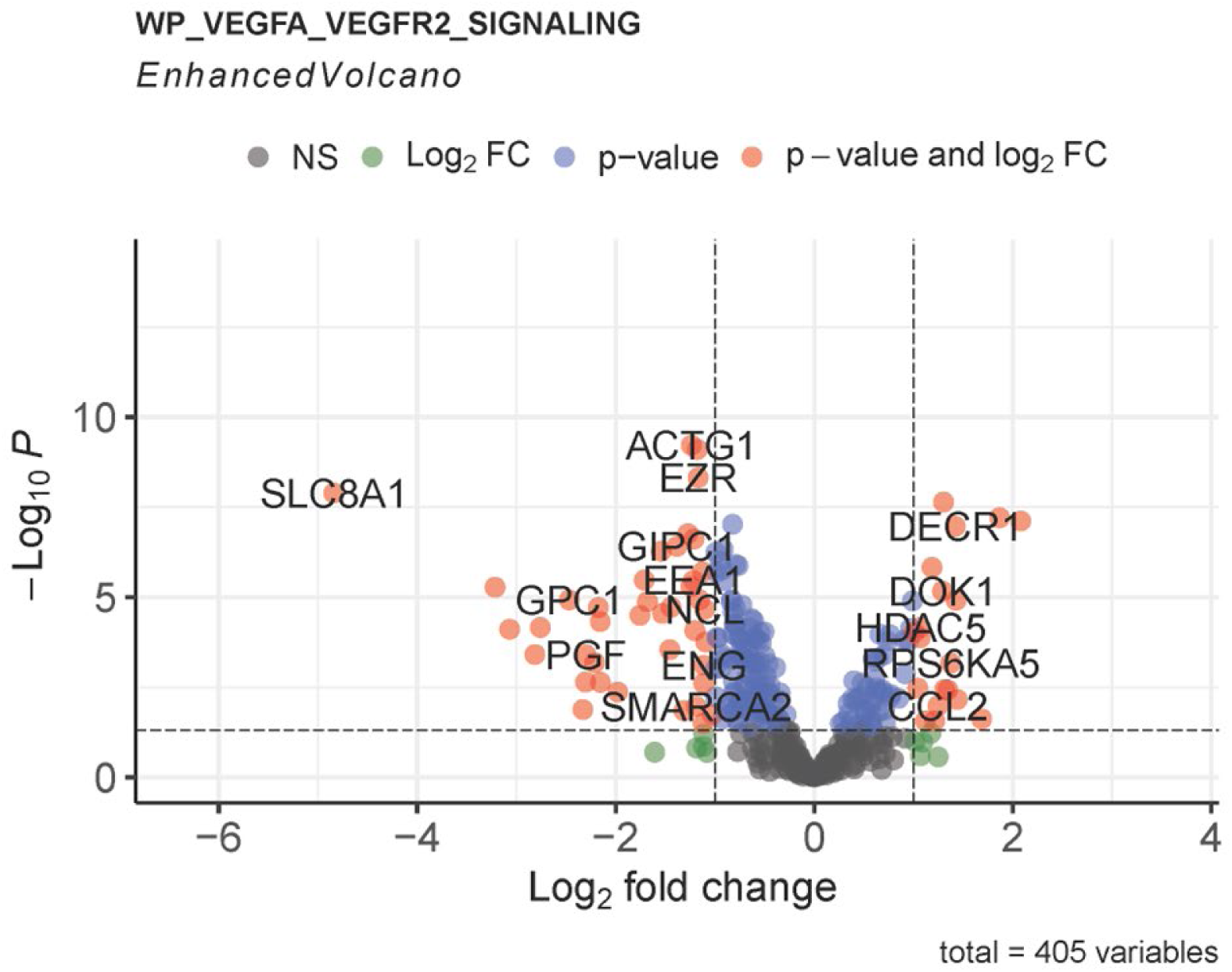
(A) VEGFA_VEGFR2 Signaling Pathway volcano plot showing a higher fraction of genes downregulated (negative fold change) in MD-1 compared to ND-1.

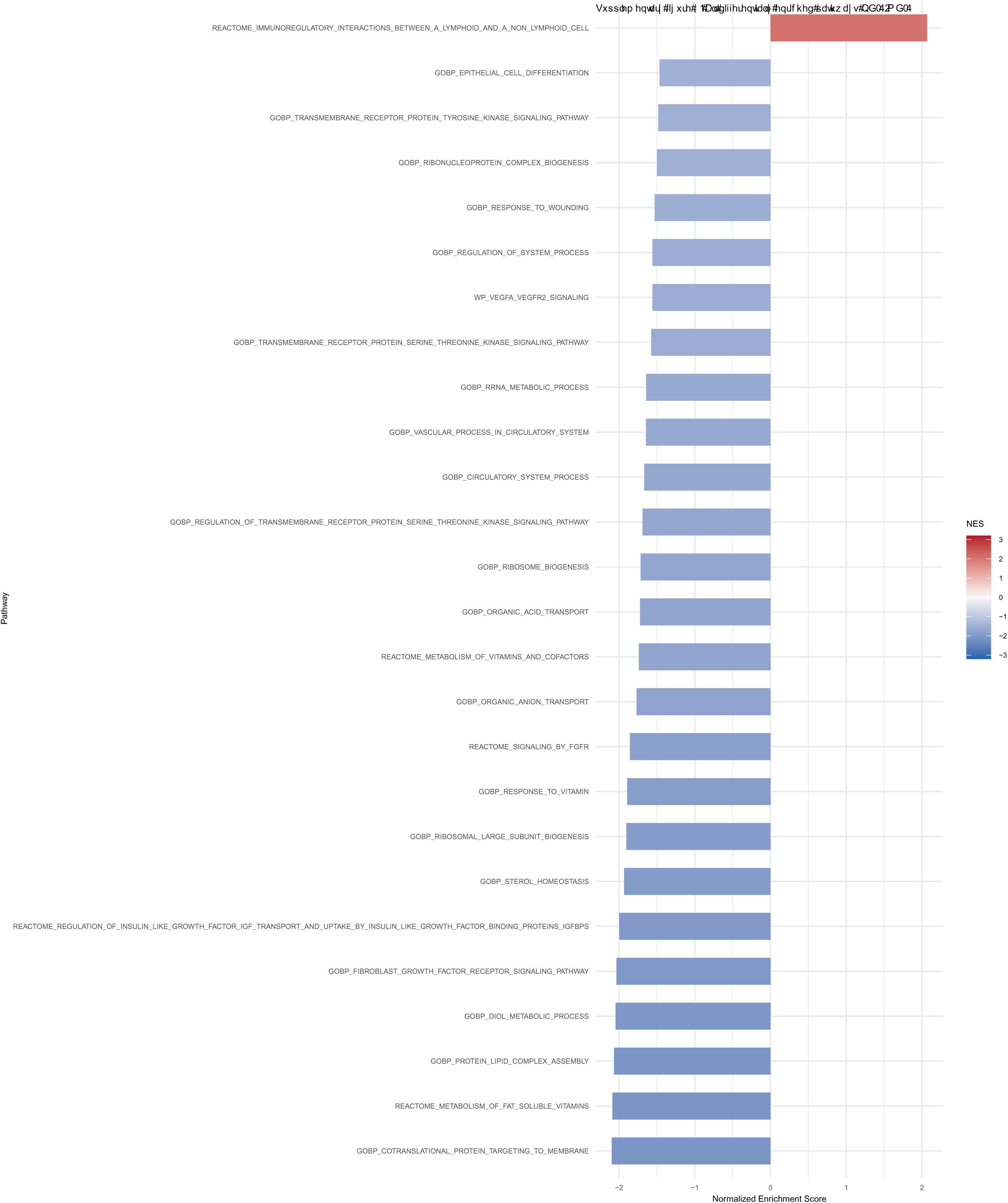

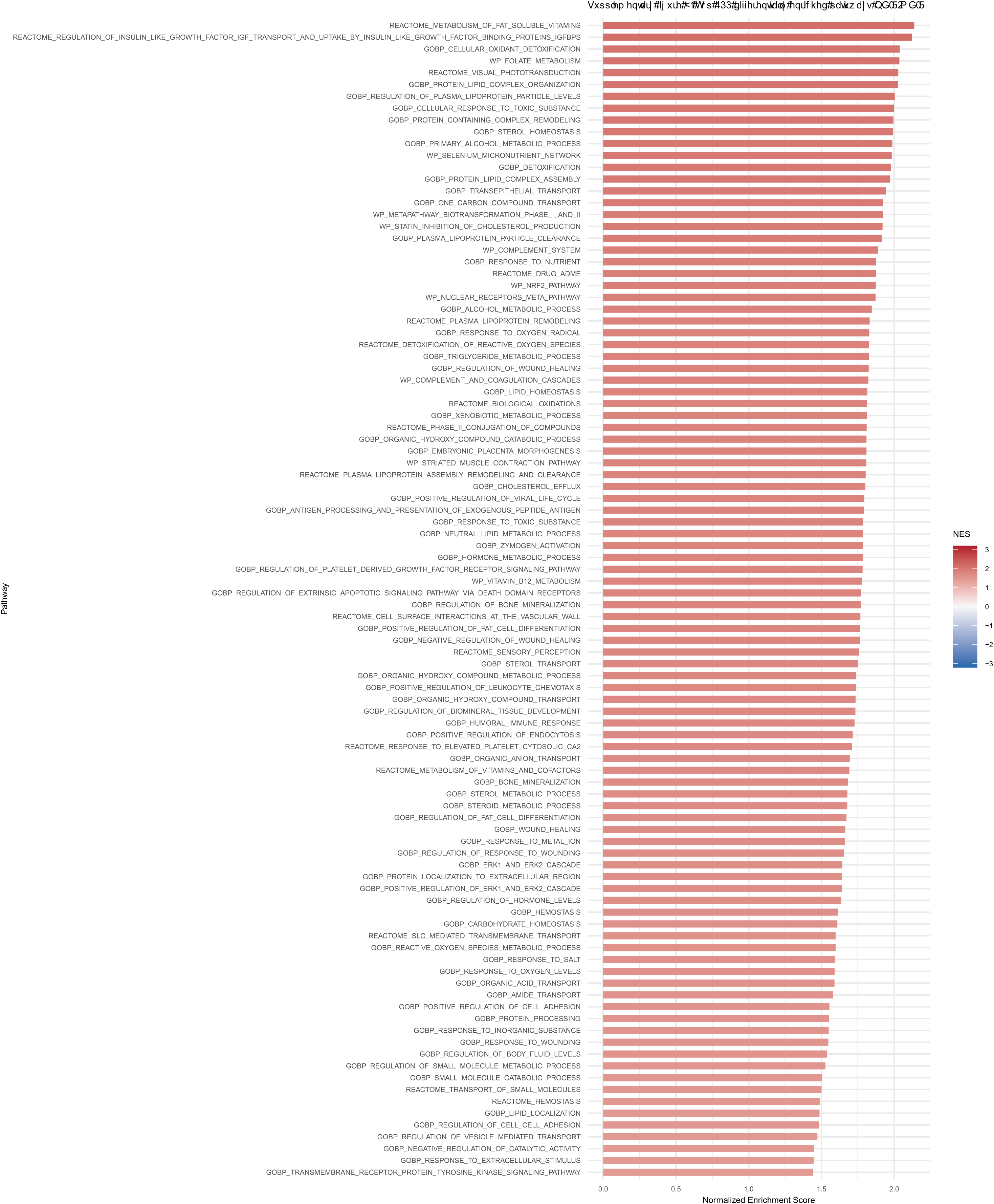

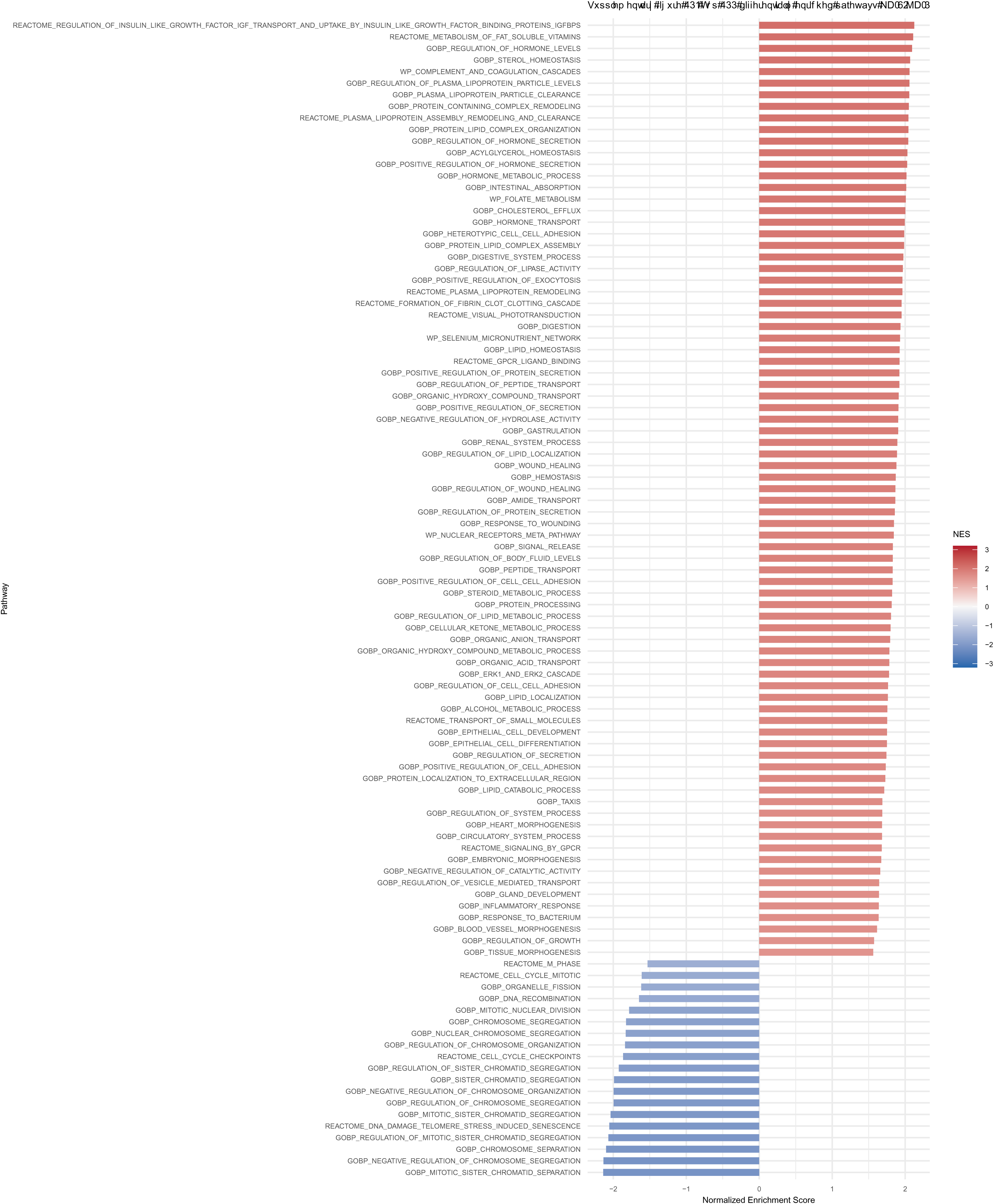

